# Kinetoplastid kinetochore proteins KKT14-KKT15 are divergent Bub1/BubR1-Bub3 proteins

**DOI:** 10.1101/2024.01.04.574194

**Authors:** Daniel Ballmer, William Carter, Jolien J. E. van Hooff, Eelco C. Tromer, Midori Ishii, Patryk Ludzia, Bungo Akiyoshi

## Abstract

Faithful transmission of genetic material is crucial for the survival of all organisms. In many eukaryotes, a feedback control mechanism called the spindle checkpoint ensures chromosome segregation fidelity by delaying cell cycle progression until all chromosomes achieve proper attachment to the mitotic spindle. Kinetochores are the macromolecular complexes that act as the interface between chromosomes and spindle microtubules. While most eukaryotes have canonical kinetochore proteins that are widely conserved, kinetoplastids such as *Trypanosoma brucei* have a seemingly unique set of kinetochore proteins including KKT1–25. Available evidence suggests that *T. brucei* lacks components of the spindle checkpoint and is unable to delay cell cycle progression in response to spindle defects. It therefore remains poorly understood how kinetoplastids regulate cell cycle progression or ensure chromosome segregation fidelity. Here, we report a crystal structure of the C-terminal domain of KKT14 from *Apiculatamorpha spiralis* and uncover that it is a pseudokinase. Its structure is most similar to the kinase domain of Bub1, a key component of the spindle checkpoint in other eukaryotes. In addition, KKT14 in *T. brucei* and other trypanosomatids has a putative ABBA motif that is present in Bub1 and its paralog BubR1. We also find that the N-terminal part of KKT14 interacts with KKT15, whose WD40 repeat beta-propeller is phylogenetically closely related to a direct interactor of Bub1/BubR1 called Bub3. Our findings indicate that KKT14-KKT15 are divergent orthologs of Bub1/BubR1-Bub3, which promote accurate chromosome segregation in trypanosomes.

## Introduction

Accurate transmission of genetic material from mother to daughter cells is essential for the survival of all organisms. Segregation of replicated chromosomes in eukaryotes is achieved by a macromolecular protein complex called the kinetochore, which links centromeric DNA to microtubules (Musacchio and Desai, 2017; McAinsh and Marston, 2022). Once all chromosomes have achieved proper attachments, the multi-subunit E3 ubiquitin ligase complex called the anaphase-promoting complex/cyclosome (APC/C) gets activated. This leads to the degradation of anaphase inhibitors securin and cyclin B, triggering sister chromatid separation and exit from mitosis (Pines, 2011; Alfieri et al., 2017).

The spindle checkpoint is a surveillance system that monitors defects in kinetochore-microtubule attachments and delays the onset of anaphase (Murray, 2011). It works by inhibiting the activity of the APC/C that is in complex with its co-activator protein Cdc20. Spindle checkpoint components include Mps1, Bub1, BubR1 (Mad3), Bub3, Mad1, and Mad2 (London and Biggins, 2014; Lara-Gonzalez et al., 2021b; McAinsh and Kops, 2023). The mitotic checkpoint complex is a potent inhibitor of APC/C^Cdc20^, which in humans consists of Mad2, Cdc20, BubR1 and Bub3 (Sudakin et al., 2001). Unattached kinetochores recruit these checkpoint proteins to catalyze the formation of the mitotic checkpoint complex (De Antoni et al., 2005). Kinetochore recruitment of spindle checkpoint proteins therefore needs to be under tight control. Bub3, a WD40 repeat domain protein, directs Bub1 and BubR1 (Mad3) to kinetochores by recognizing the Mps1-phosphorylated MELT motif of the outer kinetochore protein KNL1 (Taylor et al., 1998; London et al., 2012; Shepperd et al., 2012; Yamagishi et al., 2012; Primorac et al., 2013). Bub1 and BubR1 are paralogous proteins (Suijkerbuijk et al., 2012), which have a kinase and pseudokinase domain in its C-terminus, respectively, while Mad3, a Bub1 paralog in yeast, does not have a kinase domain. The kinase activity of Bub1 is largely dispensable for its spindle checkpoint function (Sharp-Baker and Chen, 2001; Kawashima et al., 2010). The pseudokinase domain in human BubR1 is thought to play a role in ensuring its protein stability (Suijkerbuijk et al., 2012). Bub1, BubR1, and Mad3 carry the Gle2-binding sequence (GLEBS) motif that binds Bub3 and the ABBA motif that interacts with Cdc20 (Larsen et al., 2007; Diaz-Martinez et al., 2015; Di Fiore et al., 2015; Tromer et al., 2016).

Despite its importance in ensuring accurate chromosome segregation, some organisms (e.g. yeasts, fruit flies, and human HAP1 cells) do not require the spindle checkpoint for their proliferation or development under normal conditions (Hoyt et al., 1991; Li and Murray, 1991; Burds et al., 2005; Buffin et al., 2007; Raaijmakers et al., 2018). It is thought these organisms do not require a feedback-induced mitotic delay because all chromosomes can establish proper kinetochore-microtubule attachments before the APC/C gets fully activated. Furthermore, spindle checkpoint components are apparently absent in some organisms, including *Trypanosoma brucei*, which cannot halt cell cycle progression in response to spindle defects (Ploubidou et al., 1999; Morrissette and Sibley, 2002; Vleugel et al., 2012; Markova et al., 2016; Kops et al., 2020). *T. brucei* is an experimentally tractable parasite that belongs to kinetoplastids, a class of unicellular flagellated eukaryotes that are highly divergent from traditional model eukaryotes (Figueiredo et al., 2009; Cavalier-Smith, 2010; Wheeler et al., 2019). They include the parasitic order Trypanosomatida (e.g. *T. brucei*, *Trypanosoma cruzi* and *Leishmania*), free-living Bodonida (e.g. *Bodo saltans*), both belonging to the subclass Metakinetoplastina, and Prokinetoplastina (e.g. *Apiculatamorpha spiralis*, *Papus ankaliazontas*, and *Perkinsela*) (d’Avila-Levy et al., 2015). Although kinetoplastids lack essentially all spindle checkpoint components, they have a Mad2-like protein, Cdc20, and components of the APC/C (Berriman et al., 2005; Kumar and Wang, 2005). However, in *T. brucei,* the Mad2-like protein localizes near basal bodies, not kinetochores, while Cdc20 lacks a well-conserved Mad2-interacting motif, suggesting that these proteins are unable to play a role in the canonical spindle checkpoint control (Akiyoshi and Gull, 2013; Akiyoshi, 2020). By contrast, forced stabilization of cyclin B or treatment with proteasome inhibitors cause the nucleus to arrest in metaphase (Mutomba et al., 1997; Hayashi and Akiyoshi, 2018). It is therefore conceivable that trypanosomes may possess a mechanism that intrinsically regulates the timing of nuclear division by modulating the cyclin B protein level in a Mad2-independent manner.

Like spindle checkpoint components, kinetochore proteins are widely conserved among eukaryotes (van Hooff et al., 2017; Tromer et al., 2019). However, unique kinetochore proteins called Kinetoplastid KineTochore 1–25 (KKT1–25) and KKT-Interacting Protein 1–12 (KKIP1–12) are present in *T. brucei* (Akiyoshi and Gull, 2014; Nerusheva and Akiyoshi, 2016; D’Archivio and Wickstead, 2017; Nerusheva et al., 2019; Brusini et al., 2021). These proteins are conserved among kinetoplastids but do not have a significant sequence similarity to canonical kinetochore proteins, meaning that they are attractive drug targets against diseases caused by kinetoplastid parasites (Rao et al., 2019; Saldivia et al., 2020). Understanding the structure and function of kinetoplastid kinetochore proteins also has the potential to shed light on fundamental requirements for the chromosome segregation machinery in eukaryotes.

In this study, we focus on KKT14 and KKT15, proteins of unknown functions that localize at the kinetochore from G2 until the end of anaphase in *T. brucei* (Akiyoshi and Gull, 2014). Previous bioinformatics analysis using advanced Hidden Markov Model (HMM) searches (e.g. HMMER and HHpred) of *T. brucei* KKT14 failed to identify any obvious conserved domain. KKT15 has WD40 repeats that likely form a beta-propeller, a domain found in many different proteins, including Bub3, Cdc20, and mRNA export factor Rae1/Gle2 (Jain and Pandey, 2018). Here we discover that KKT14 has a pseudokinase domain in its C-terminus, which is most similar to the kinase domain of Bub1. We also identify a putative ABBA motif in KKT14. The N-terminal part of KKT14 interacts with KKT15, which we suggest to be a Bub3 ortholog. These results reveal that kinetoplastids possess divergent Bub1/BubR1 and Bub3 proteins.

## Results

### Crystal structure of KKT14 C-terminal domain reveals similarity to Bub1

To gain insights into the function and evolutionary origin of KKT14, we aimed to obtain its high-resolution structural information. By screening four kinetoplastid species (*T. brucei*, *T. cruzi*, *Paratrypanosoma confusum*, and *Apiculatamorpha spiralis*), we succeeded in determining a 2.2 Å resolution crystal structure for the C-terminal domain of the KKT14 protein from the prokinetoplastid *A. spiralis* (clone PhF-6) (Figure 1A, Table 1 and Table S1). *A. spiralis* KKT14^365−640^ crystallized with two molecules in an asymmetric unit. Both molecules were essentially identical except for minor variations in flexible loops.

**Figure 1.**
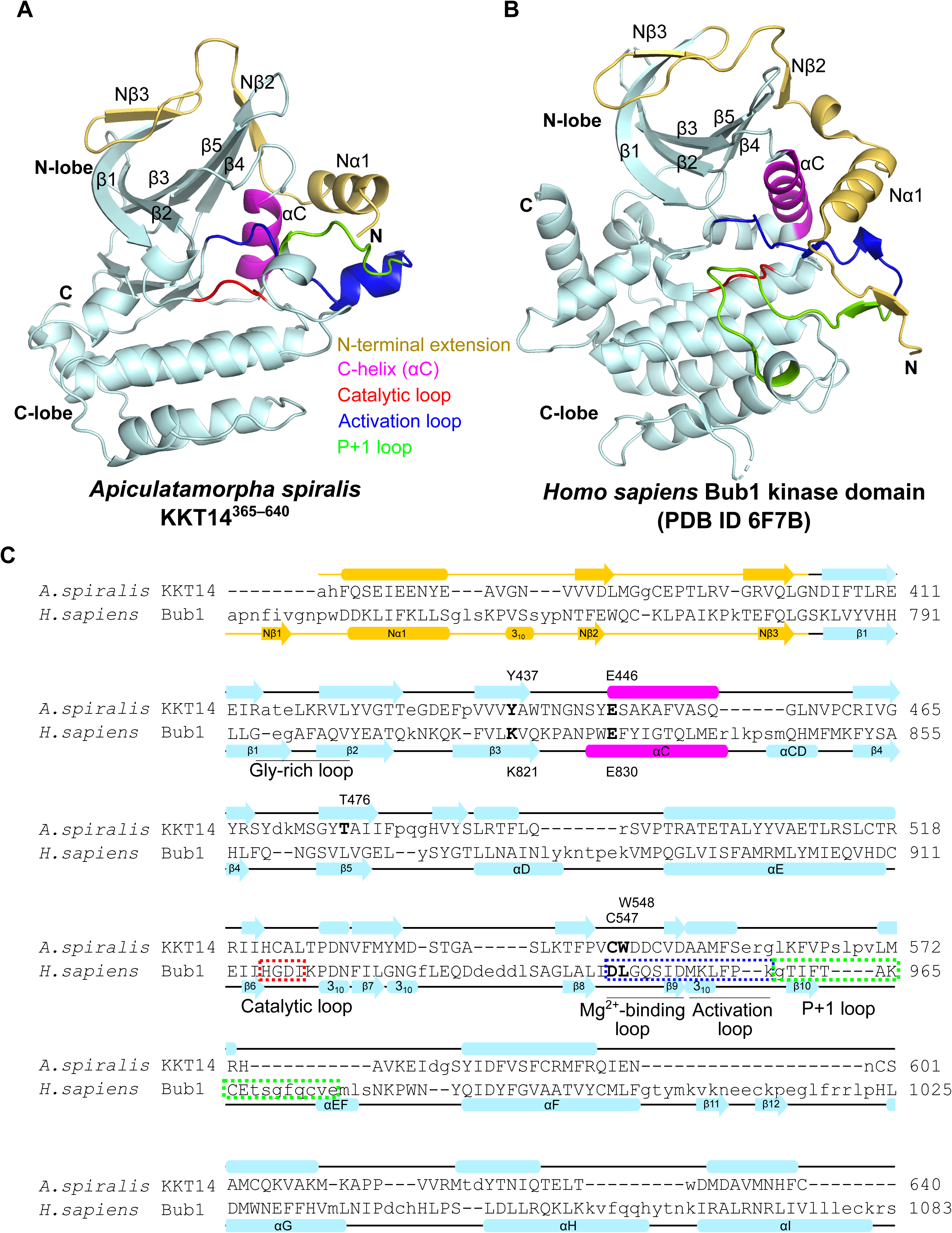
Crystal structure of *A. spiralis* KKT14 reveals similarity to the Bub1 kinase domain. (A, B) Cartoon representation of *A. spiralis* KKT14^365–640^ (A) and human Bub1 kinase domain (PDB accession 6F7B (Siemeister et al., 2019)) (B). The fold nomenclature of the N-terminal extension and the kinase domain of Bub1 is based on (Kang et al., 2008). (C) Structure-based pairwise alignment of *A. spiralis* KKT14 and human Bub1 kinase domain based on the DALI search output. Structurally equivalent residues are in uppercase, while structurally non-equivalent residues (e.g. in loops) are in lowercase. Secondary structures were assigned using DSSP (Kabsch and Sander, 1983).

**Table 1:** Data collection and refinement statistics for *Apiculatamorpha spiralis* KKT14^365−640^.

| **Data collection** |  |
| --- | --- |
| Beamline | Diamond Light Source I03 |
| Wavelength (Å) | 0.9763 |
| Space group (Z) | *P* 1 2<sub>1</sub> 1 |
| Unit cell | 49.76 Å 87.58 Å 71.88 Å<br>90° 93.61° 90° |
| Resolution range (Å) | 49.66–1.95 (1.95–1.99) |
| Unique reflections | 41803 (2243) |
| Completeness (%) | 93.3 (100.0) |
| Multiplicity | 6.7 (6.7) |
| *I*/*σ* | 6.9 (0.7) |
| *R*<sub>meas</sub> | 0.188 (2.640) |
| CC1/2 | 1.0 (0.6) |
| Wilson B-factor (Å<sup>2</sup>) | 30.14 |
| **Refinement** |  |
| No. reflections | 28797 (1146) |
| *R*<sub>work</sub> | 0.22 (0.36) |
| *R*<sub>free</sub> | 0.24 (0.40) |
| Number of atoms | 4571 |
| Protein | 4308 |
| Solvent | 263 |
| RMS bonds (Å) | 0.009 |
| RMS angles (°) | 1.23 |
| Ramachandran favored (%) | 96.13 |
| Ramachandran allowed (%) | 3.87 |
| Ramachandran outliers (%) | 0.00 |
| Average B-factor (Å<sup>2</sup>) | 39.00 |

Values in parentheses correspond to the highest resolution shell

Interestingly, a structural homology search using the distance-matrix alignment (DALI) server (Holm et al., 2023) revealed similarity to a protein kinase fold with an N-lobe and C-lobe (Figure 1A and Table S2). The N-lobe contains a five-stranded β-sheet, a helix termed the C-helix (αC), and loops that correspond to the catalytic loop and activation loop, while the C-lobe comprises a bundle of α-helices (Figure 1A). Although most protein kinases share a similar fold (Endicott et al., 2012), we found that the most similar structure of *A. spiralis* KKT14^365−640^ in the PDB database was the Bub1 kinase domain (Figure 1B). Human Bub1 was the top hit in the DALI search with a Z-score of 15.5, while the next best hit was the MST3 kinase (Z-score 13.0) (Table S2). We obtained a similar result for an AlphaFold2-predicted structure of *T. brucei* KKT14^358–685^ (Figure S1A), showing a Z-score of 15.9 for human Bub1 and 12.9 for the next best hit, the PAK3 kinase (Table S2). Moreover, searches with Foldseek against the AlphaFold2-predicted structure database (van Kempen et al., 2023) produced congruent results (Methods). The higher structural similarity of KKT14 to the kinase domain of Bub1 rather than other kinases is due to an extended N-terminal helix that is present in Bub1 and KKT14 (Figure 1A–C, Table S2) (Kang et al., 2008; Lin et al., 2014; Breit et al., 2015; Huang et al., 2019; Siemeister et al., 2019). These results show that the C-terminal domain of KKT14 has a kinase fold with the most similar structure being the Bub1 kinase domain, raising a possibility that KKT14 is a Bub1/BubR1 ortholog.

Besides a kinase/pseudokinase domain, Bub1 and BubR1 have various conserved domains and motifs (Diaz-Martinez et al., 2015; Di Fiore et al., 2015; Davey and Morgan, 2016; Tromer et al., 2016). We identify a putative ABBA motif (consensus: Fx[ILV][FHY]x[DE]) in KKT14, which is highly conserved among trypanosomatids (Figure 2), as well as KEN boxes in some kinetoplastids (Figure S2). In contrast, other domains such as a TPR, CDI or a KARD domain were not found. The presence of an ABBA motif and a C-terminal Bub1-like kinase fold strongly supports the possibility that KKT14 is a divergent Bub1-like protein.

**Figure 2.**
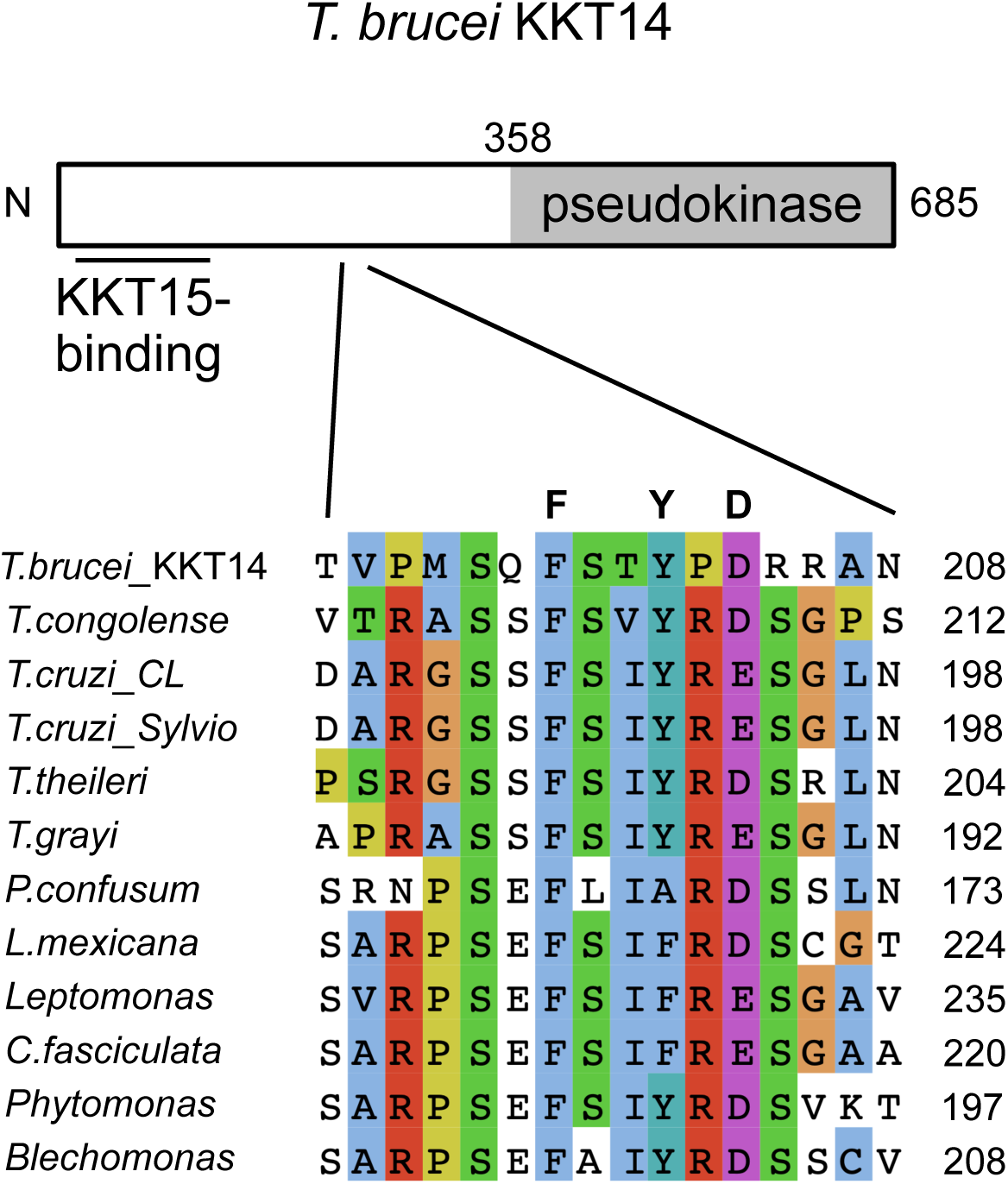
KKT14 has a putative ABBA motif. Schematic of *T. brucei* KKT14 protein and a multiple sequence alignment showing a putative ABBA motif (consensus Fx[ILV][FHY]x[DE] based on (Di Fiore et al., 2015)) conserved in trypanosomatid KKT14 proteins.

### KKT14 C-terminal domain is an inactive pseudokinase

Despite the structural similarity, however, a structure-based sequence alignment of KKT14 with Bub1 shows a lack of conserved residues that play key roles in catalytically-active protein kinases (Figure 1C). Most notably, none of KKT14 orthologs in kinetoplastids has a conserved β3 lysine (K821 in Bub1) (Figure 3A), whose mutation results in an inactive kinase (Tang et al., 2004; Bayliss et al., 2012). In addition, KKT14 does not appear to have the Gly-rich motif (GxGxxG in most kinases, which is essential for stabilizing ATP phosphates during catalysis), and the HRD motif in the catalytic loop (usually His-Arg-Asp: HGD in Bub1) is HGN in *T. brucei* and HCA in *A. spiralis* (Figure 3A). Furthermore, the DFG motif (usually Asp-Phe-Gly: DLG in Bub1), which is required for Mg^2+^ coordination, is HWE in *T. brucei* and CWD in *A. spiralis* KKT14 (Figure 3A). Given that even a single amino acid change of key residues in these motifs results in inactive kinases (Kwon et al., 2019), these findings strongly suggest that the C-terminal domain of KKT14 is a catalytically-inactive pseudokinase. Consistent with this possibility, KKT14 purified from trypanosomes did not have any detectable auto-phosphorylation activity, while the KKT3 kinase did (Figure 3B).

**Figure 3.**
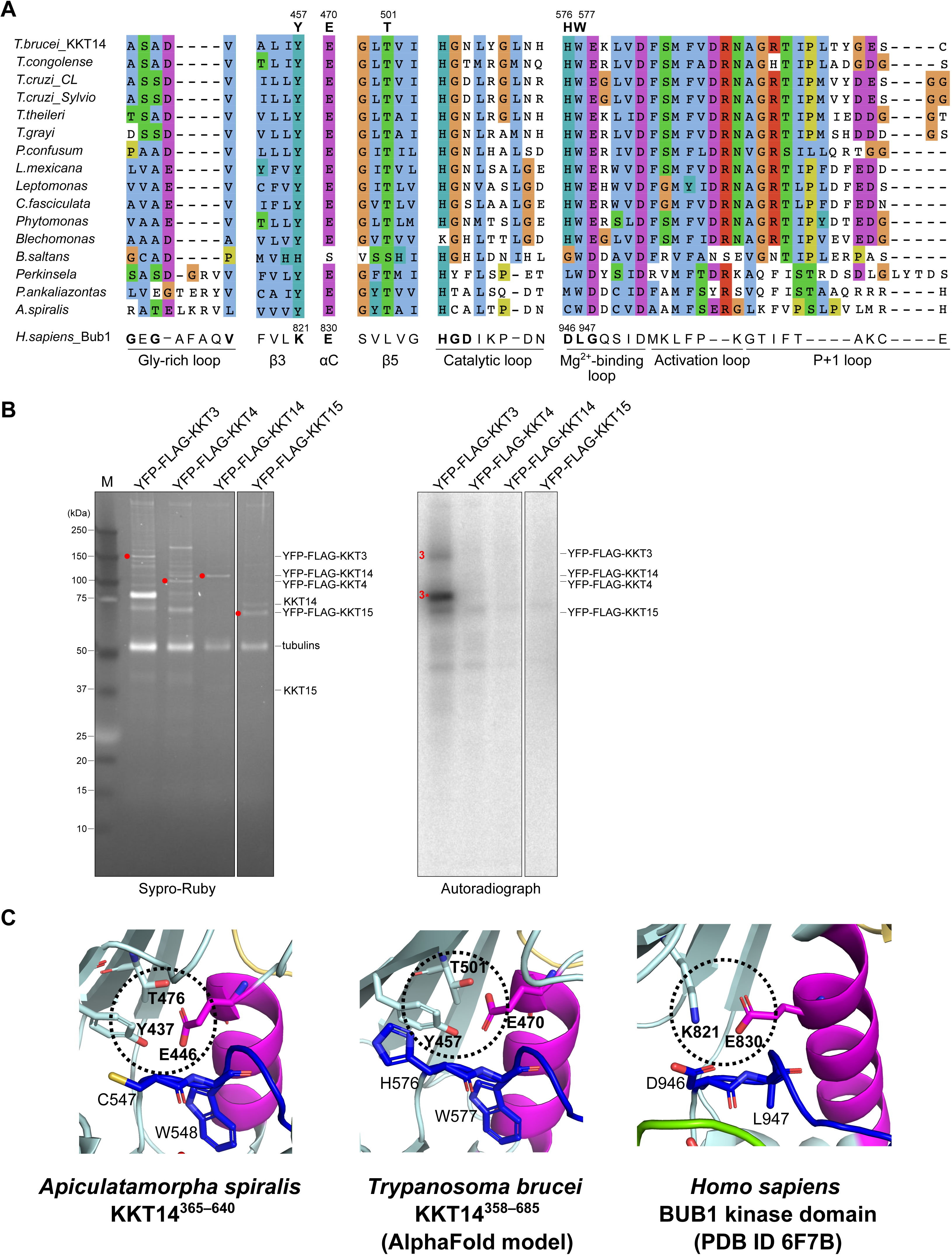
KKT14 lacks key residues that are present in active protein kinases. (A) Multiple sequence alignment of kinetoplastid KKT14 sequences highlighting the regions that correspond to the key parts of the Bub1 kinase domain. Note that T501 and W577 in *T. brucei* are conserved among kinetoplastids. (B) Lack of detectable auto-phosphorylation activity for KKT14. Indicated proteins were immunoprecipitated from trypanosomes using FLAG antibodies and eluted with FLAG peptides. The left panel shows a Sypro-Ruby stained SDS-PAGE gel (red circles indicate FLAG-tagged proteins), while the right panel shows phosphorylation detected by autoradiography. A degradation product of YFP-FLAG-KKT3 is indicated by 3*. (C) The C-helix has an “in” conformation in the *A. spiralis* KKT14 crystal structure and AlphaFold2-predicted *T. brucei* KKT14 structure (Supplementary Dataset S1). Note that E446 in the C-helix is in close proximity with Y437 of β3 and T476 of β5 in *A. spiralis* (E470, T501, and Y457 in *T. brucei* respectively), while E830 in the C-helix forms a salt bridge with the conserved β3 lysine (K821) in Bub1.

Although KKT14 appears to be an inactive pseudokinase, features of active protein kinase conformations are found in its structure. In the active conformation of protein kinases, the DFG motif and the C-helix have an “in” conformation, where the conserved phenylalanine or leucine of the DFG motif (L947 in Bub1) points out of the active site and the aspartic acid (D946 in Bub1) faces the ATP-binding site (Figure 3C) (Bayliss et al., 2012). In addition, the β3 lysine (K821 in Bub1) forms a salt bridge with the glutamate (E830 in Bub1) in the C-helix (Figure 3C). In the crystal structure of *A. spiralis* KKT14, W548 (the phenylalanine equivalent of the DFG motif, which is CWD in *A. spiralis* KKT14), strictly conserved among kinetoplastids, faces away from the active site. Even though KKT14 in almost all kinetoplastids has the conserved glutamate in the C-helix (E446 in *A. spiralis*), the position of β3 lysine has tyrosine (Y437 in *A. spiralis*). Nonetheless, the C-helix sits close to β3, mediated by apparent interactions among Y437, E446, and T476 in β5. Similar interactions were observed in the AlphaFold2-predicted structure of *T. brucei* KKT14 (Figure 3C), and the threonine is strictly conserved among kinetoplastids (T501 in *T. brucei*) (Figure 3A). We also note that the positions of the N-terminal extension, C-helix, catalytic loop, and Mg^2+^-binding loop are very similar in between the crystal structure of *A. spiralis* KKT14 and the AlphaFold2-predicted structure of *T. brucei* KKT14 (Figure S1), despite a limited similarity between the KKT14 sequences of the two species (20.1% identical, 31.5% similar). These findings suggest that the catalytically-inactive pseudokinase domain of KKT14 takes an active-like conformation of a kinase fold.

### KKT15 is a divergent Bub3 protein

Our previous sequence similarity searches were unsuccessful in finding homology between *T. brucei* KKT14 and Bub1/BubR1. Using HHpred (Zimmermann et al., 2018) with a sensitive alignment of KKT14 proteins from an extended kinetoplastid dataset (Table S3), we were now able to observe a link to human Bub1, albeit with a non-significant E-value (Table S4). Due to the low level of sequence similarities, conducting a phylogenetic analysis was not feasible. In contrast, searches with a KKT15 alignment retrieved many WD40 domain proteins with highly significant E-values and had Bub3 as its best hit (Table S4). The WD40 beta-propeller is a domain present in Bub3 and many other proteins (Jain and Pandey, 2018). To investigate to which WD40 group KKT15 might belong, we conducted a phylogenetic analysis using two multiple sequence alignments of WD40 proteins (Methods). In both cases, KKT15 orthologs cluster with Bub3 and Rae1, which are known to be closely related to one another (Tromer et al., 2019). One alignment yields all KKT15 orthologs as sister to Bub3 and Rae1 (Figure 4 and Figure S3A), whereas the other places only a particular subset, those of *B. saltans*, *A. spiralis* and *P. ankaliazontas*, next to Bub3 and Rae1 (Figure S3B), suggesting that trypanosomatid KKT15 sequences differ more from Bub3/Rae1 than those of bodonids and prokinetoplastids. Importantly, KKT15 proteins do not cluster within Bub3 in either tree, so we cannot unequivocally designate them as Bub3 orthologs. However, the fact that kinetoplastids have clear Rae1 orthologs (Figure S3A,B) (Billington et al., 2023), in combination with its kinetochore localization and interaction with KKT14 (see below), analogous to Bub1/BubR1-Bub3, prompts us to propose that KKT15 is a Bub3 ortholog.

**Figure 4.**
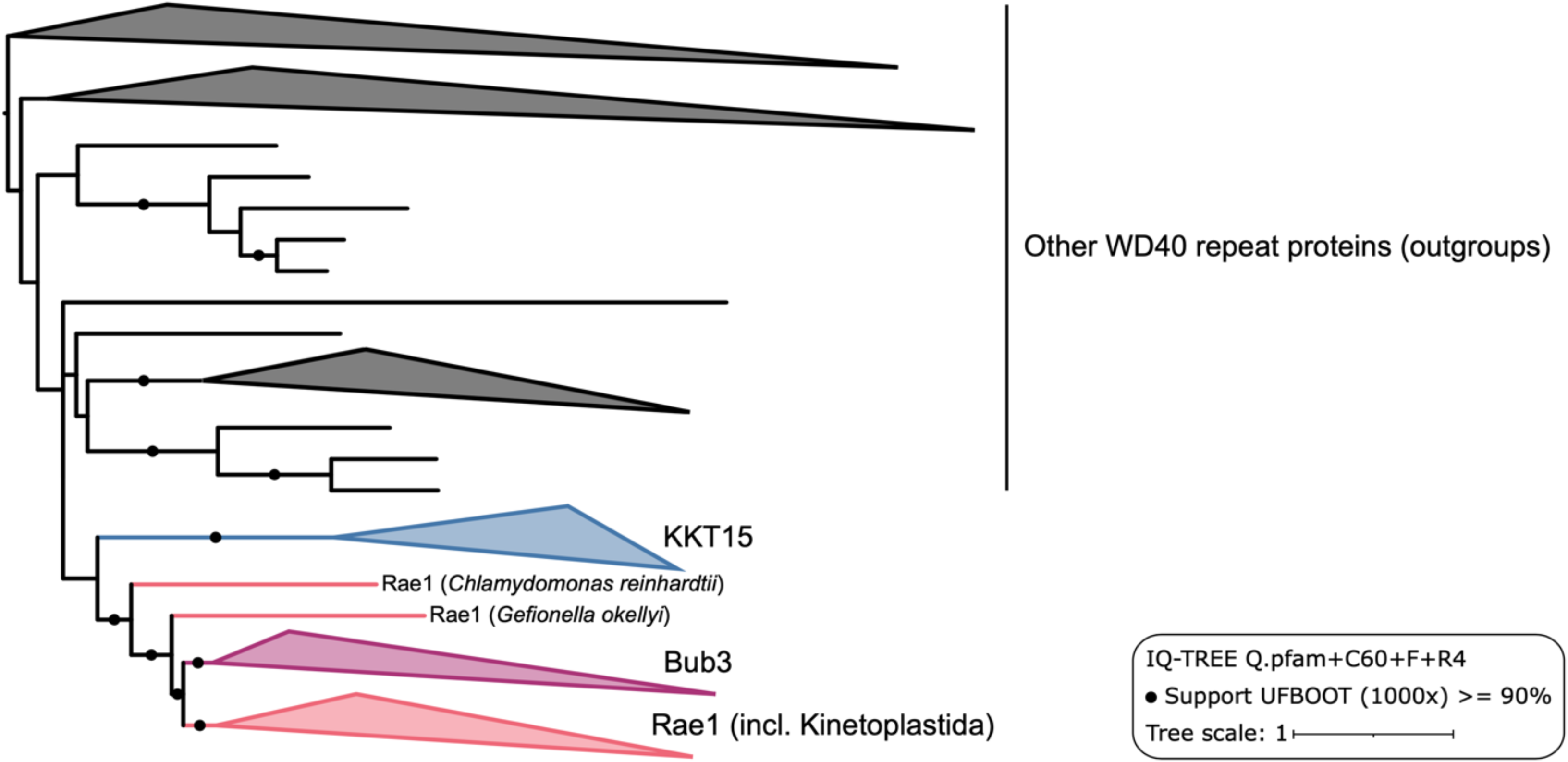
Phylogeny of KKT15 and related WD40 repeat proteins. Subtree of the full phylogeny presented in Supplementary Figure S3A. Note that in the alignment approach applied here, KKT15 proteins were prompted to form a single group (see Methods).

### KKT14 interacts with KKT15

KKT14 and KKT15 localize at kinetochores from G2 to anaphase (Akiyoshi and Gull, 2014). This localization pattern differs from the rest of transiently-localized kinetoplastid kinetochore proteins that start to localize at kinetochores from S phase, suggesting that KKT14 and KKT15 may directly interact with each other. This possibility is supported by our mass spectrometry analysis (Figure 5A). Although KKT14 does not appear to have a GLEBS motif present in the N-terminal region of Bub1/BubR1 that interacts with Bub3 (Larsen and Harrison, 2004; Larsen et al., 2007; Primorac et al., 2013), AlphaFold2 predicted an interaction between the N-terminal region of KKT14 and KKT15 (Figure 5B, 5C). The region of KKT14 predicted to interact with KKT15 is well conserved among trypanosomatids (residues 2–111 in *T. brucei*) (Figure S2). Interestingly, the confidence score (predicted local distance difference test: pLDDT) for this region improved when predicted as a KKT14 – KKT15 complex, compared to KKT14 alone (Figure 5D), implying that this region of KKT14 is more likely to form a secondary structure (α helices) in the complex. These results strongly support the idea that kinetoplastid kinetochore proteins KKT14 and KKT15 are divergent Bub1/BubR1 and Bub3 proteins, although they might have adopted a distinct interaction mode.

**Figure 5.**
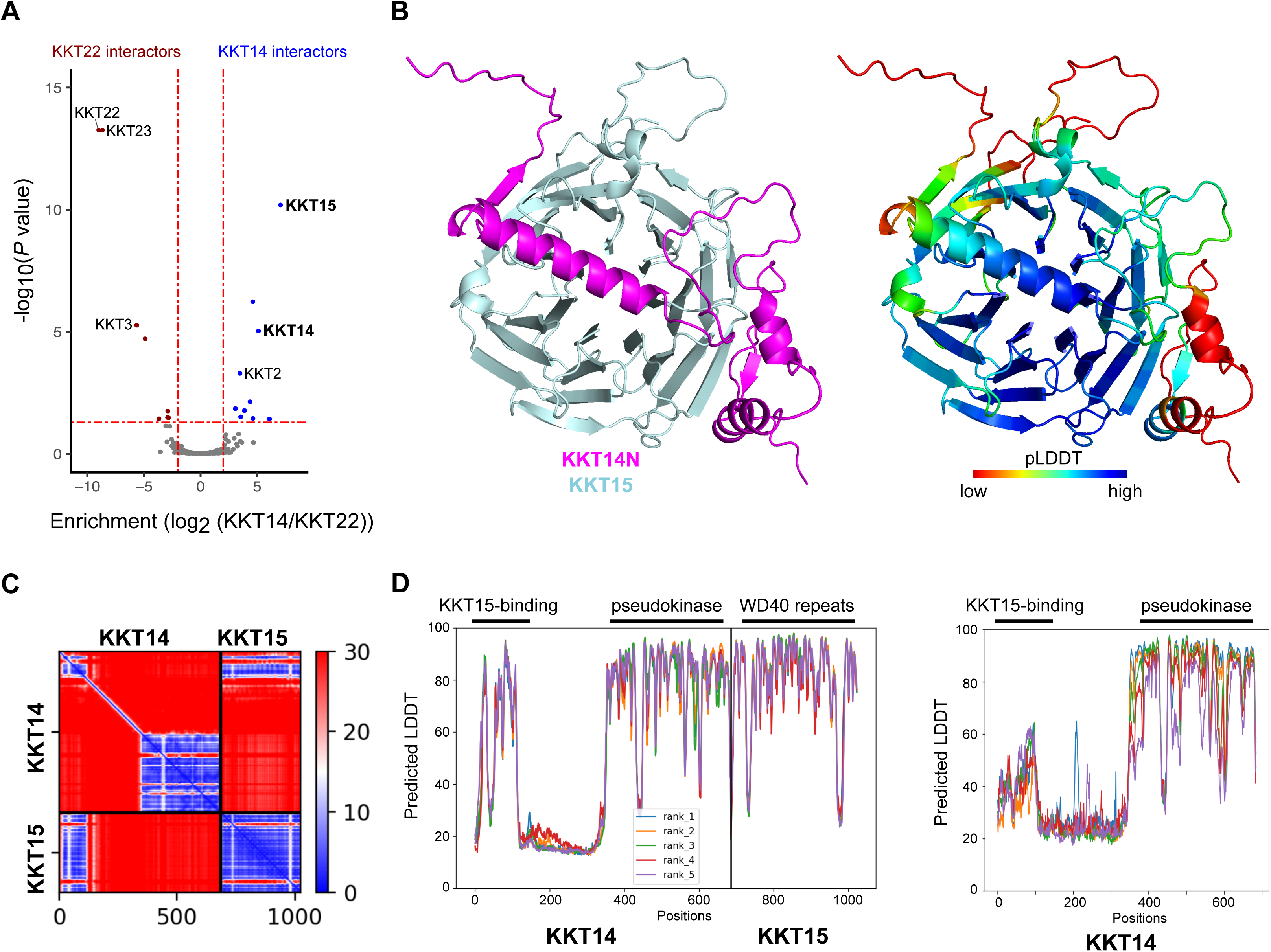
KKT14 is predicted to interact with KKT15 directly. (A) A volcano plot showing relative enrichment and significance values between the immunoprecipitates of YFP-KKT14 and YFP-KKT22 (n=4 each). KKT22 was used as a comparison, which mainly co-purifies with KKT23 and KKT3 (Nerusheva et al., 2019). See Table S5 for all proteins identified by mass spectrometry. (B) AlphaFold2 predictions of the KKT14^2–125^-KKT15 complex in cartoon representation (Supplementary Dataset S2). (C) The Predicted Aligned Error (PAE) plots for the rank 1 model for KKT14-KKT15, predicting interactions via the N-terminal part of KKT14. (D) Predicted local distance difference test (pLDDT) plots for KKT14-KKT15 (left) and KKT14 (right). AlphaFold2-predicted models are provided in the Supplementary Dataset S3 and S4.

To better characterize KKT14, we ectopically expressed its fragments in trypanosomes. We found that KKT14N^2–357^ localized at kinetochores from G2 to anaphase, while KKT14C^358–685^ only had diffuse nuclear signals (Figure 6A). Immunoprecipitation of these fragments revealed that KKT14N co-purified with many kinetochore proteins, including KKT15 (Figure 6B and Table S5). Furthermore, LacO/LacI-based tethering experiments show that KKT14N, not KKT14C, was able to recruit KKT15 to an ectopic locus in vivo (Figure 6C). These results suggest that the N-terminal region of KKT14 interacts with KKT15, as predicted by AlphaFold2 (Figure 5B).

**Figure 6.**
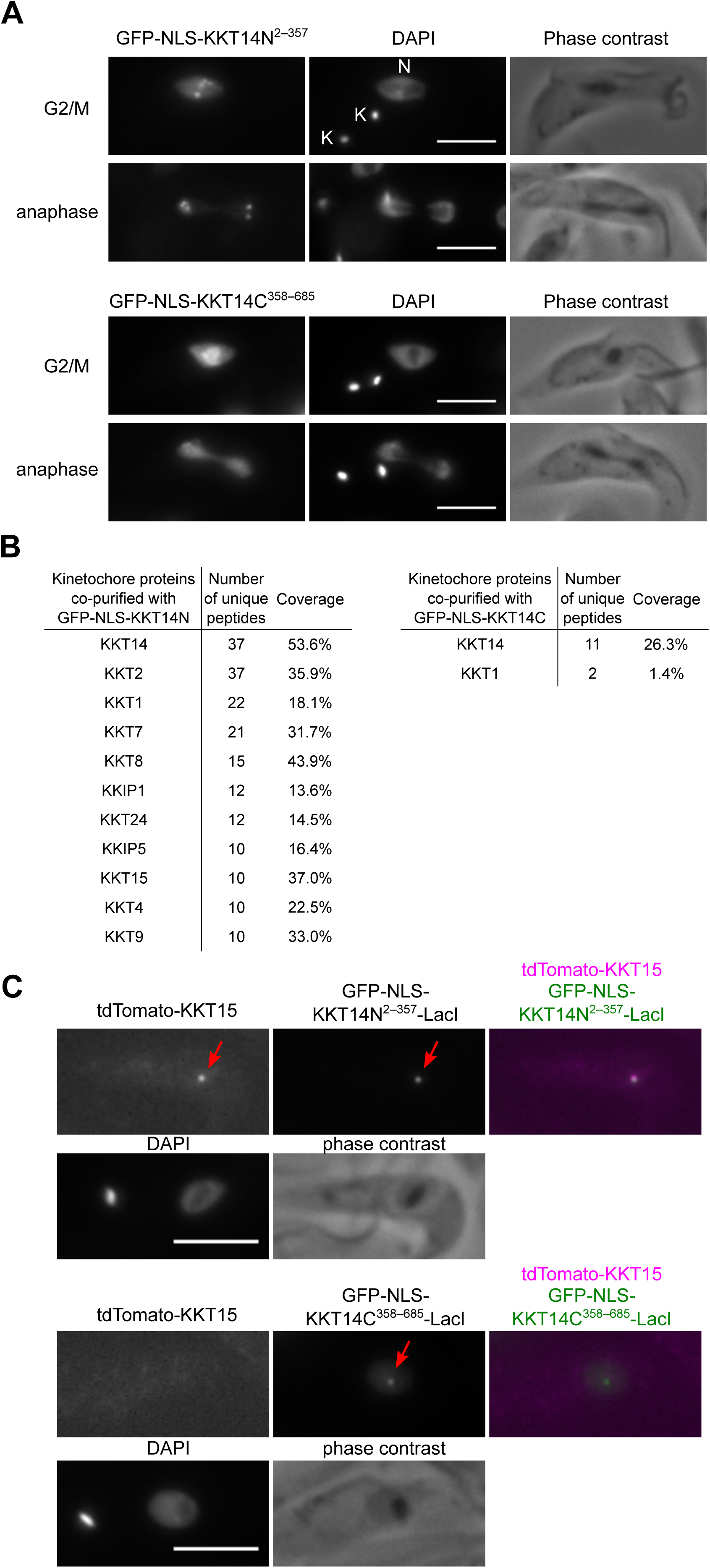
N-terminal region of KKT14 binds KKT15. (A) Ectopically expressed GFP-KKT14N^2–357^ localizes at kinetochores, while GFP-KKT14C^358–685^ does not. K and N stand for the kinetoplast (mitochondrial DNA) and nucleus, respectively. Cell cycle stages of individual cells were determined based on the number of K and N as described previously (Woodward and Gull, 1990; Siegel et al., 2008). The GFP fusion proteins were expressed in trypanosomes using 10 ng/mL doxycycline for 1 day and fixed for microscopy. Cell lines: BAP2386, BAP2387. (B) Immunoprecipitation/mass spectrometry analysis shows that GFP-KKT14N co-purifies with many kinetochore proteins, while GFP-KKT14C does not. Immunoprecipitation was carried out using cells expressing the GFP fusion proteins using 10 ng/mL doxycycline for 1 day. See Table S5 for all proteins identified by mass spectrometry. (C) KKT14N^2–357^ is sufficient to recruit KKT15 in trypanosomes. Recruitment of tdTomato-KKT15 was observed in 100% or 0% of 1K1N (G1) cells that have GFP-KKT14N^2–357^-LacI or GFP-KKT14C^358–685^-LacI dots respectively (n=10 each). The GFP fusion proteins were expressed in trypanosomes using 10 ng/mL doxycycline for 1 day. Cell lines: BAP2655, BAP2656. Bars, 5 µm.

### KKT14 and KKT15 are required for accurate chromosome segregation

We next performed an RNAi-mediated knockdown of KKT14 and KKT15 to assess their function for chromosome segregation. The RNAi construct for KKT14 is previously described (Marcianò et al., 2021), while that for KKT15 was established in this study (Figure 7A–C). We found that kinetochore localization of KKT14 and KKT15 are mutually co-dependent (Figure 7D–G), further supporting the notion that they form a complex. Although KKT14 depletion caused severe growth defects (Figure 7H) (Marcianò et al., 2021), we failed to find obvious cell cycle profile changes at 8 or 16 hr after induction of KKT14 RNAi, apart from a moderate increase in anaphase cells (Figure 7I). In contrast, we observed lagging kinetochores in almost all anaphase cells even at 8 hr post-induction (Figure 7J, K). These results show that KKT14 is essential for accurate chromosome segregation and cell growth.

**Figure 7.**
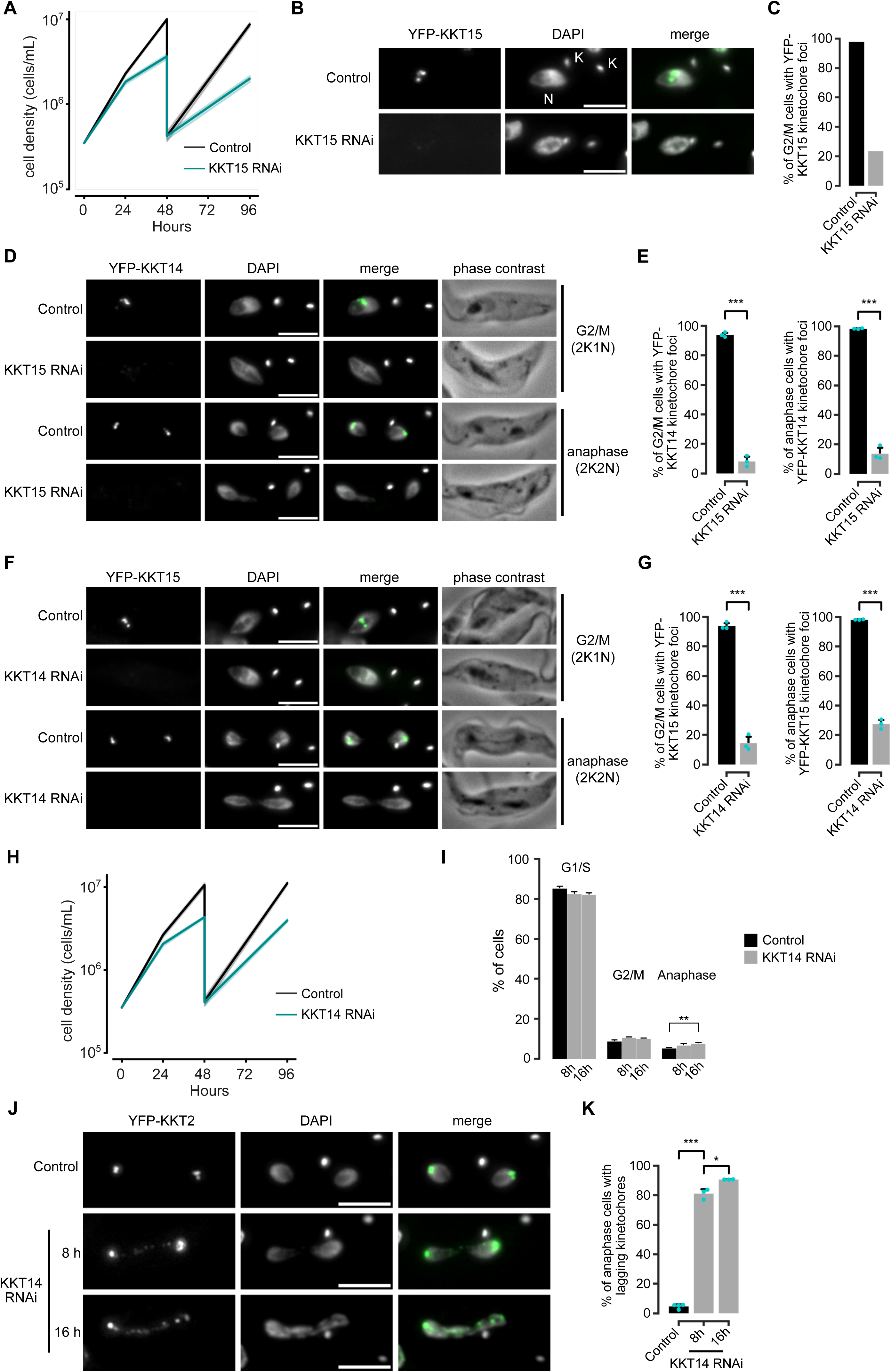
KKT14 is essential for accurate chromosome segregation in trypanosomes. (A) Growth curve upon RNAi-mediated knockdown of KKT15 using an RNAi construct against its 3’ UTR. Data are presented as the mean ± SD of three replicates. RNAi was induced with 1 μg/mL doxycycline and cultures were diluted at day 2. Cell line: BAP2533. (B) and (C) Validation of KKT15 knockdown. RNAi was induced with 1 μg/mL doxycycline and cultures were diluted at day 2. K and N stand for the kinetoplast and nucleus, respectively. Cell line: BAP2535. At least 130 cells per condition were quantified. (D) Representative fluorescence micrographs showing the localization of YFP-KKT14 upon RNAi-mediated knockdown of KKT15 in G2/M (2K1N) and anaphase (2K2N) cells. RNAi was induced with 1 μg/mL doxycycline for 24 h. Cell line: BAP2533. (E) Quantification of 2K1N and 2K2N that have kinetochore-like dots of YFP-KKT14 upon RNAi-mediated depletion of KKT15. All graphs depict the means (bar) ± SD of three replicates (shown as dots). (F) Representative fluorescence micrographs showing the localization of YFP-KKT15 upon RNAi-mediated knockdown of KKT14 in 2K1N and 2K2N cells. RNAi was induced with 1 μg/mL doxycycline for 24 h. Cell line: BAP2534. (G) Quantification of 2K1N and 2K2N that have kinetochore-like dots of YFP-KKT15 upon RNAi-mediated depletion of KKT14. All graphs depict the means (bar) ± SD of three replicates (shown as dots). A minimum of 50 cells per replicate were quantified in each condition. (H) Growth curve upon RNAi-mediated knockdown of KKT14. Data are presented as the mean ± SD of three replicates. RNAi was induced with 1 μg/mL doxycycline and cultures were diluted at day 2. Cell line: BAP2534. (I) Cell cycle profile upon knockdown of KKT14. RNAi was induced with 1 μg/mL doxycycline and cells were fixed at 8 h or 16 h. All graphs depict the means (bar) ± SD of at three replicates. A minimum of 700 cells per replicate were quantified. Cell line: BAP680. (J) Representative fluorescence micrographs showing lagging kinetochores (marked by tdTomato-KKT2) in anaphase cells upon KKT14 knockdown at 8 and 16 h post-induction. RNAi was induced with 1 μg/mL doxycycline. Cell line: BAP680. (K) Quantification of lagging kinetochores in anaphase cells upon KKT14 knockdown. All graphs depict the means (bar) ± SD of three replicates (shown as dots). A minimum of 50 cells per replicate were quantified in each condition. * P < 0.05, ** P ≤ 0.01, *** P ≤ 0.001 (two-sided, unpaired t-test).

## Discussion

In this study, we propose that the kinetoplastid kinetochore proteins KKT14 and KKT15 are divergent Bub1/BubR1 and Bub3 orthologs, respectively. The discovery of a kinase fold in KKT14 was surprising because our previous sequence-based approach or a previous study that comprehensively cataloged pseudokinases failed to identify KKT14 as a pseudokinase (Kwon et al., 2019). In other words, KKT14’s pseudokinase domain is highly divergent from other kinases or pseudokinases. It is therefore remarkable that KKT14 retains significant structural similarities to the kinase domain of Bub1, including its N-terminal extension. In human Bub1, this N-terminal extension acts as a “minicyclin” by making extensive contacts with the N-lobe of the kinase domain and thereby promoting an active conformation (Kang et al., 2008). Similarly, in both the crystal structure of *A. spiralis* KKT14 and AlphaFold2-predicted structure of *T. brucei* KKT14, the N-terminal extension makes extensive contacts with the N-lobe, which may stabilize the pseudokinase structure. Although the function of the KKT14 pseudokinase domain remains unclear, it is striking that all known KKT14 orthologs in kinetoplastids conserved it.

In contrast to KKT14, KKT15 has a readily discernible, common protein fold, namely a WD40 repeat beta-propeller. Our phylogenetic analysis places KKT15 close to Bub3. Taking into account also its interaction with the divergent Bub1/BubR1 protein KKT14, we propose that KKT15 is a Bub3 ortholog. Yet, its sequence is quite divergent from Bub3, particularly in trypanosomatids. In humans, Bub3 localizes at kinetochores by recognizing the phosphorylated KNL1 protein, which is not found in kinetoplastids. Furthermore, it remains unclear whether KKT15 binds a phosphorylated peptide because the regions of Bub3 that bind KNL1’s phosphorylated MELT motif (Primorac et al., 2013) are not well conserved in KKT15 (Figure S4 and S5). It therefore remains unknown how the KKT14-KKT15 complex is recruited to kinetochores. Depletion of KKT2 disrupts KKT14 localization, suggesting that KKT14 and KKT15 are downstream of KKT2 (Marcianò et al., 2021). Although our mass spectrometry data support a possibility that KKT14 and/or KKT15 directly interact with KKT2, AlphaFold2 fails to predict interactions between them. It will be important to identify direct interaction partners for KKT14 and KKT15 to reveal how these proteins function at kinetoplastid kinetochores.

In human and *C. elegans*, some of the ABBA motifs in Bub1/BubR1 function by promoting kinetochore localization of Cdc20 and contribute to the strength of the checkpoint (Di Fiore et al., 2015; Vleugel et al., 2015; Kim et al., 2017; Lara-Gonzalez et al., 2021a; Piano et al., 2021). In trypanosomes, kinetochore localization of Cdc20 has not been reported ((Billington et al., 2023) and our unpublished data), and it remains unclear if, how, when, and where the ABBA motif of KKT14 may regulate the activity of Cdc20 and contribute to the cell cycle control. Addressing these questions will be key to understanding how KKT14 and KKT15 contribute to accurate chromosome segregation in trypanosomes that lack a canonical spindle checkpoint.

## Materials and Methods

### Trypanosomes and microscopy

All trypanosome cell lines used in this study were derived from *T. brucei* SmOxP927 procyclic form cells (TREU 927/4 expressing T7 RNA polymerase and the tetracycline repressor to allow inducible expression) (Poon et al., 2012) and are listed in Table S1. Cells were grown at 28°C in SDM-79 medium supplemented with 10% (vol/vol) heat-inactivated fetal calf serum, 7.5 µg/mL hemin (Brun and Schönenberger, 1979), and appropriate drugs. Endogenous YFP tagging was performed using the pEnT5-Y vector (Kelly et al., 2007) or a PCR-based method (Dean et al., 2015). Endogenous 3FLAG-6HIS-YFP tagging was performed using pBA106 (Hayashi and Akiyoshi, 2018), while endogenous tdTomato tagging was performed using pBA892 (Ishii and Akiyoshi, 2020). LacO-LacI tethering experiments were performed as described previously using the LacO array inserted at the rDNA locus (Landeira and Navarro, 2007; Ishii and Akiyoshi, 2020). Inducible expression of GFP-NLS fusion and GFP-NLS-LacI fusion proteins was carried out using pBA310 (Nerusheva and Akiyoshi, 2016) and pBA795 (Ishii and Akiyoshi, 2020), respectively. Cell growth was monitored using a CASY cell counter (Roche). Expression of GFP fusion proteins and RNAi were induced with doxycycline at a final concentration of 10 ng/mL and 1 µg/mL, respectively. All plasmids were linearized by *Not*I and transfected into trypanosomes by electroporation. Transfected cells were selected by the addition of 30 μg/mL G418 (Sigma), 25 μg/mL hygromycin (Sigma), 5 μg/mL phleomycin (Sigma), or 10 μg/mL blasticidin S (Insight biotechnology).

### Immunoprecipitation

For each experiment, 400-mL cultures of asynchronously growing cells (unless otherwise indicated) were grown to ∼1×10^7^ cells/mL and harvested. Ectopic expression of GFP-tagged KKT14N^2–357^ and KKT14C^358–685^ in *T. brucei* was induced with 10 ng/mL doxycycline for 24 hr. YFP-KKT14 and YFP-KKT22 were expressed from the endogenous locus. Where indicated, 10 µM MG132 (to arrest cells prior to anaphase) or 2 µM 1NM-PP1 (to inhibit the AUK1 kinase activity) were added for 4 hr prior to harvesting the cells. Note that there was no noticeable change in the amount of co-purifying proteins in these conditions, so we pooled all results for the volcano plot analysis (Figure 5A and Table S5). Immunoprecipitation of GFP/YFP-tagged proteins was performed with anti-GFP antibodies (11814460001, Roche) using a method we previously described (Ishii and Akiyoshi, 2020). 3FLAG-6HIS-YFP-tagged proteins expressed from the endogenous locus were immunoprecipitated using anti-FLAG M2 antibodies (F3165, Sigma) (Unnikrishnan et al., 2012) and eluted with 0.5 mg/mL 3×FLAG peptide (F4799, Sigma) in BH0.15 (25 mM HEPES pH 8.0, 2 mM MgCl_2_, 0.1 mM EDTA pH 8.0, 0.5 mM EGTA pH 8.0, 1% NP-40, 150 mM KCl, and 15% glycerol) supplemented with protease inhibitors (10 μg/mL leupeptin, 10 μg/mL pepstatin, 10 μg/mL E-64, and 0.2 mM PMSF) and phosphatase inhibitors (1 mM sodium pyrophosphate, 2 mM Na-β-glycerophosphate, 0.1 mM Na_3_VO_4_, 5 mM NaF, and 100 nM microcystin-LR) with agitation for 25 min at room temperature. FLAG eluates were run on an SDS-PAGE gel, which was stained with Sypro-Ruby (Thermo Fisher).

### In vitro kinase assay

To examine auto-phosphorylation activities in the immunoprecipitated 3FLAG-6HIS-YFP-KKT3/4/14/15 samples, 10 µL of FLAG eluates were mixed with 2.5 µL of 10× kinase buffer (500 mM Tris-HCl pH 7.4, 10 mM DTT, 250 mM β-glycerophosphate, 50 mM MgCl_2_, 50 μCi [^32^P] ATP, and 100 μM ATP) in 25 µL volumes. The mixture was incubated at 30°C for 30 min, and the reaction was stopped by the addition of the LDS sample buffer (Thermo Fisher). The samples were run on an SDS-PAGE gel and stained with Coomassie Brilliant Blue R-250 (Bio-Rad) (not shown), which was subsequently dried and used for autoradiography using a phosphorimager screen. The signal was detected by an FLA 7000 scanner (GE Healthcare).

### Mass spectrometry

Reduction of disulfide bridges in cysteine-containing proteins was performed with 10 mM DTT dissolved in 50 mM HEPES, pH 8.5 (56°C, 30 min). Reduced cysteines were alkylated with 20 mM 2-chloroacetamide dissolved in 50 mM HEPES, pH 8.5 (room temperature, in the dark, 30 min). Mass spectrometry samples were prepared using the SP3 protocol (Hughes et al., 2019), and trypsin (Promega) was added in the 1:50 enzyme to protein ratio for overnight digestion at 37°C. Next day, peptide recovery was done by collecting supernatant on magnet and combining with second elution of beads with 50 mM HEPES, pH 8.5. For a further sample clean up, an OASIS HLB µElution Plate (Waters) was used. The samples were dissolved in 10 µL of reconstitution buffer (96:4 water: acetonitrile, 1% formic acid and analyzed by LC-MS/MS using QExactive (Thermo Fisher) in the proteomics core facility at EMBL Heidelberg (https://www.embl.org/groups/proteomics/). Peptides were identified by searching tandem mass spectrometry spectra against the *T. brucei* protein database with MaxQuant (version 2.0.1) with carbamidomethyl cysteine set as a fixed modification and oxidization (Met), phosphorylation (Ser, Thr, and/or Tyr), and acetylation (N-term and Lys) set as variable modifications. Up to two missed cleavages were allowed. The first peptide tolerance was set to 10 ppm (protein FDR 1%). Proteins identified with at least two peptides were considered significant and reported in Table S5. All raw mass spectrometry files and the custom database file used in this study have been deposited to the ProteomeXchange Consortium via the PRIDE partner repository (Perez-Riverol et al., 2022; Deutsch et al., 2023) with the dataset identifier PXD047806.

Differential enrichment analysis of YFP-KKT14 vs YFP-KKT22 was performed on iBAQ values using the DEP package in R (Zhang et al., 2018) (Table S5). Reverse hits and contaminants were removed, and results were filtered for proteins that were identified in all replicates of at least one condition. The data was background corrected and normalized by variance stabilizing transformation (vsn). Missing values were imputed using the k-nearest neighbor approach (knn). Potential interactors were determined using t-tests, with threshold values set to lfc = 2 and alpha = 0.05. The volcano plot shown was constructed using the EnhancedVolcano package (Blighe, 2023).

### Expression and purification of *Apiculatamorpha spiralis* KKT14C

To make pBA2356 (6HIS-KKT14^365–640^ from *Apiculatamorpha spiralis* (clone PhF-6)), the DNA was amplified from BAG142 (a synthetic DNA that encodes *Apiculatamorpha spiralis* KKT14, codon optimized for expression in *E. coli*) with primers BA3187/BA3188 and cloned into RSFDuet-1 using *Bam*HI/*Eco*RI sites with the NEBuilder HiFi DNA Assembly kit (NEB) (Table S1). *E. coli* BL21(DE3) cells were transformed with ∼100 ng of plasmid DNA (pBA2356) and inoculated into 50 mL of 2×TY medium containing 50 μg/mL kanamycin and grown overnight at 37°C. The next morning, 6 L of 2×TY medium with 50 μg/mL of kanamycin was warmed at 37°C, and 5 mL of the overnight culture was inoculated into each liter. Cells were grown at 37°C with shaking (200 rpm) until the OD_600_ reached ∼0.6. Protein expression was induced with 0.2 mM IPTG for 16 hr at 20°C. Cells were spun down at 3,400 g at 4°C and resuspended in 200 mL of lysis buffer (50 mM sodium phosphate, pH 7.5, 500 mM NaCl, and 10% glycerol) supplemented with protease inhibitors (20 μg/mL leupeptin, 20 μg/mL pepstatin, 20 μg/mL E-64 and 0.4 mM PMSF), benzonase nuclease (500 U per 1 liter culture), and 0.5 mM TCEP. All subsequent steps were performed at 4°C. Bacterial cultures were mechanically disrupted using a French press (1 passage at 20,000 psi) and the soluble fraction was separated by centrifugation at 48,000 g for 30 min. Supernatants were loaded on 5 mL of TALON beads (Takara Bio) pre-equilibrated with the lysis buffer. Next, the beads were washed with 300 mL of the lysis buffer with 0.5 mM TCEP, and proteins were eluted with 50 mM sodium phosphate pH 7.5, 500 mM NaCl, 10% glycerol, 250 mM imidazole and 0.5 mM TCEP. To cleave off the His-tag, samples were incubated with TEV protease in 1:50 w/w ratio overnight while being buffer-exchanged into 25 mM sodium phosphate, 250 mM NaCl, 5% glycerol, 5 mM imidazole, and 0.5 mM TCEP by dialysis. To increase the sample purity and remove the His-tag, samples were re-loaded on TALON beads pre-equilibrated with the dialysis buffer and the flow-through was collected. Next, the sample was concentrated using 10-kD MW Amicon concentrator (Millipore), and loaded on Superdex 75 16/600 (GE Healthcare) columns to further purify and buffer exchange into 25 mM HEPES pH 7.5, 150 mM NaCl with 0.5 mM TCEP. Fractions containing the protein of interest were pooled, concentrated to 15.1 mg/mL using a 10-kD MW Amicon concentrator (Millipore), and flash-frozen in liquid nitrogen for –80°C storage.

### Crystallization trials and structural determination

All crystals were obtained in sitting drop vapor diffusion experiments in 96-well plates, using drops of overall volume 200 nL, mixing protein and mother liquor in a 1:1 v/v ratio. Crystals of *A. spiralis* KKT14^365−640^ (15.1 mg/mL) were grown at 4°C in MIDAS HT-96 B1 solution (Molecular Dimensions) containing 0.1 M sodium formate and 20% w/v SOKALAN CP 45. Crystals were briefly transferred into mother liquor prepared with addition of 25% glycerol prior to flash-cooling by plunging into liquid nitrogen. Data collection and model building X-ray diffraction data from *A. spiralis* KKT14^365−640^ were carried out at the i03 beamline at the Diamond Light Source (Harwell, UK). The structure was solved using the AlphaFold2 predicted structure of *A. spiralis* KKT14^398−640^ as a model with a molecular replacement software, PHASER (McCoy, 2017) followed by initial model building with BUCCANEER (Cowtan, 2006). The data were scaled to 2.2 Å based on I/Iσ parameters (I/Iσ value of 2.0 was used as a threshold). Further manual model building and refinement were completed iteratively using COOT (Emsley et al., 2010) and PHENIX (Liebschner et al., 2019). All images were made with PyMOL (version 2.5.2, Schrödinger). Protein coordinates have been deposited in the RCSB protein data bank with the accession number 8QOH.

### Bioinformatic analysis of KKT14 and KKT15

The protein sequences for KKT14 and KKT15 were retrieved from the TriTryp database (Aslett et al., 2010) or published studies (Butenko et al., 2020; Tikhonenkov et al., 2021). Searches for their homologous proteins were done using BLAST in the TriTryp database (Aslett et al., 2010) or manual searches using hmmsearch (HMMER version 3.0) on predicted proteomes using manually prepared hmm profiles (Eddy, 1998). Multiple sequence alignments were performed with MAFFT (L-INS-i method, version 7) (Katoh et al., 2019) and visualized with the clustalx coloring scheme in Jalview (version 2.11) (Waterhouse et al., 2009). The pairwise sequence identity and similarity between *A. spiralis* KKT14^365−640^ and *T. brucei* KKT14^358–685^ were calculated using EMBOSS Needle (Madeira et al., 2022). Structures and interactions were predicted with AlphaFold2-Multimer-v2.3.1 (Jumper et al., 2021; Evans et al., 2022) through ColabFold version 1.5.3 using MMseqs2 with 24 recycles (UniRef+Environmental) (Mirdita et al., 2022). The rank 1 model, PAE, and pLDDT plots for each prediction are provided in the supplemental material (Dataset S1–4). In all cases, similar results were obtained for five predictions. All structure figures were made using PyMOL version 2.5.2 (Schrödinger, LLC). The following command was used to map pLDDT score onto the AlphaFold2-predicted structure models: spectrum b, rainbow_rev, maximum=100, minimum=50. Foldseek searches were carried out using the web server against AlphaFold2-predicted structure databases covering UniProt50, Swiss-Prot, and Proteome (version 4, Mode 3Di/AA) (https://search.foldseek.com/search) (Varadi et al., 2022; van Kempen et al., 2023).

### Phylogenetic analysis of KKT15 and related WD40 repeat-containing proteins

To conduct a phylogenetic analysis to determine to which WD40 repeat proteins KKT15 proteins are closely related, we selected the 30 best EggNOG KOG/COG hits resulting from online HHpred (Zimmermann et al., 2018; Gabler et al., 2020) of our best, most sensitive KKT15 multiple sequence alignment. The latter was obtained by iterative profile HMM searches combined with phylogenetic analysis among a local database of euglenozoa and some other eukaryotic protein sequences, as well as a local eukaryote-wide dataset (de Potter et al., 2023) (Table S3). The trusted list of KKT15 orthologs across these euglenozoa was aligned using MAFFT (v.7.505, option L-INS-i, alignment and profile HMM are included in Supplementary Dataset S5) (Katoh et al., 2019) and submitted to online HHpred to search in its COG_KOG_v1.0 database. Among these 30 best hits, there were only KOG families, and one duplicate, which we removed (Table S4). We collected the corresponding profile HMMs of these KOGs from EggNOG (v.5.0) (Huerta-Cepas et al., 2019) and added our initial KKT15 HMM. We then executed hmmscan (hmmer.org) of these HMMs versus a subsampled (49 diverse eukaryotes) version of our local eukaryotic database, applying an E-value cut-off of 1×10^-5^. We assigned the retrieved eukaryotic proteins to their respective KOG (or KKT15), and appended the original euglenozoan KKT15 orthologs to that family as well. We subsequently, again for all KOGs/KKT15, used hmmsearch among the assigned hits, in order to be able to retrieve the 100 best hits per KOG/KKT15, and gathered of these 100 hits only the domain that was hit by the HMM (the putative WD40 repeat). For Rae1 (KOG0647), we also included the hit regions of some additional Discoba ortholog candidates, because these were not part of the 49 species selection but potentially informative in placing KKT15. Except for KKT15 and these additional Discoba sequences, we subjected the KOG sequence selections to CD-hit (v4.8.1, identity cut-off 70%) (Fu et al., 2012), in order to facilitate the phylogenetic analysis and interpretation. We then removed all sequences across all KOGs that were shorter than 50 amino acids (i.e., this was not applied to KKT15). Each KOG was separately aligned using MAFFT (L-INS-i).

We then employed two different strategies to combine all sequences into a single alignment, resulting in two different phylogenies (referred to as tree 1 and tree 2, found in Supplementary Figure S3A and B, respectively). For tree 1, we similarly aligned KKT15, and subsequently used MAFFT’s –merge option to combine all individual alignments into a single alignment (parameters: –-localpair –-maxiterate 100 –-merge). We also had identified five additional, potentially close KKT15 homologs of different species, which cannot be evidently classified as a particular WD40 repeat protein. We added them to the alignment using MAFFT option –add (parameters: –-maxiterate 1000 –-add). For tree 2, we used MAFFT –merge (parameters: –-localpair –-maxiterate 100 –merge) on just the KOG’s individual alignments, and subsequently used MAFFT –add (parameters: –-maxiterate 1000 –add) to add KKT15 orthologs and the sequences of unknown identity to it. The alignments for tree 1 and tree 2 hence differ in the way the KKT15 sequences are aligned: either first among one another (tree 1), or only through adding the sequences to the already existing (merged) alignment of the KOGs. The first approach forces the KKT15 sequences to be monophyletic in the tree, while the second approach does not. We trimmed both alignments using trimAl (Capella-Gutiérrez et al., 2009) (v1.4.rev15, option –gappyout) and removed sequences with >85% gaps. We used the alignments to infer a maximum likelihood phylogeny with IQ-TREE (version 2.0.3) (Minh et al., 2020), applying an evolutionary model selected by ModelFinder (Kalyaanamoorthy et al., 2017), also allowing for complex mixture (C-series) models to be selected. Branch support was estimated through ultrafast bootstraps (1000 replicates) (Hoang et al., 2018). The tree with the highest likelihood, either the maximum likelihood tree or the consensus tree, was selected for visualization in iTOL (Letunic and Bork, 2021). The phylogenies were initially rooted on a well-supported clade with a relatively long branch, which was not closely associated with Rae1, Bub3 or KKT15 (KOG2111). The phylogenies were annotated using, if available, the name of the human protein belonging to each KOG. If not available, the budding yeast protein name was used. The multiple sequence alignments and raw IQ-TREE output can be found in Supplementary Dataset S5. The full, uncollapsed phylogenies can be inspected on iTOL: https://itol.embl.de/tree/62145194227274281702966749 (tree 1, associated with Supplementary Figure S3A) and https://itol.embl.de/tree/62145194227421141702968325 (tree 2, associated with Supplementary Figure S3B). Note that alongside KKT15, we performed a similar online HHpred search for a refined KKT14 multiple sequence alignment, the results of which are also reported in Table S4. The alignments used as input for these searches can be found in Supplementary Dataset S6.

## Acknowledgments

We thank the crystallography facility manager Edward Lowe, Micron Advanced Bioimaging Unit, and the proteomics core facility at EMBL, especially Mandy Rettel and Jennifer Schwarz, for their support. We also thank Miguel Navarro (Instituto de Parasitologıá y Biomedicina López-Neyra, Consejo Superior de Investigaciones Cientificas, Spain) for providing the pMig75 and pMig96 plasmids, and Sam Dean (Warwick Medical School, University of Warwick, UK) for pPOT plasmids. D. Ballmer was supported by the Berrow Foundation. J. J. E van Hooff was supported by a Veni Fellowship from the Dutch Research Council (NWO). E. C. Tromer was supported by a Veni Fellowship from the Dutch Research Council (NWO, VI.Veni.202.223). P. Ludzia was supported by the Boehringer Ingelheim Fonds. B. Akiyoshi was supported by a Wellcome Trust Senior Research Fellowship (grant 210622/Z/18/Z) and a Wellcome Discovery Award (grant 227243/Z/23/Z). The authors declare no competing financial interests.

## Rights retention

This research was funded in whole, or in part, by the Wellcome Trust [210622/A/18/Z and 227243/Z/23/Z]. For the purpose of open access, the author has applied a CC BY public copyright licence to any Author Accepted Manuscript version arising from this submission.

## Author contributions

D. Ballmer characterized the RNAi phenotype of KKT14 and KKT15 and performed immunoprecipitation and mass spectrometry of YFP-KKT14/KKT22. W. Carter and P. Ludzia solved and analyzed the crystal structure of *A. spiralis* KKT14. J. J. E van Hooff and E. C. Tromer performed sequence and phylogenetic analysis of KKT15/Bub3. M. Ishii established KKT14 knockdown methods. B. Akiyoshi performed kinase assays, LacO/LacI tethering experiments, AlphaFold2 analyses, and wrote the manuscript with input from all authors.

## Supplemental material

### Supplementary Tables (Excel files)

**Table S1.** List of trypanosome strains, plasmids, primers, and synthetic DNA used in this study

**Table S2.** DALI search results for *A. spiralis* KKT14 crystal structure and AlphaFold2-predicted *T. brucei* KKT14 structure

**Table S3**. Datasets of the predicted proteomes of a diversity of Discoba (sheet ‘Discoba’) and eukaryotes (sheet ‘Eukaryota’), including the type of data, their sources and BUSCO completeness scores. The sheets also contain the field ‘Abbreviation’, which contains the first part of the protein identifiers for each respective species, as they are found in the files in Supplementary Dataset.

**Table S4**. Search results of KKT15 and KKT14 alignments using online HHpred (Zimmermann et al., 2018). For each of KKT15 and KKT14, two searches were performed: one against the EggNOG database, and the other against a subset of model organisms (*Homo sapiens*, *Saccharomyces cerevisiae*, *Arabidopsis thaliana* and *Naegleria gruberi*). The top 30 of the results is included in this table. The results of KKT15 against EggNOG were used as input for the phylogenetic analysis of KKT15, including a variety of WD40 repeat proteins.

**Table S5.** List of proteins identified by mass spectrometry. Immunoprecipitation of endogenously tagged YFP-KKT14 and YFP-KKT22 and exogenously expressed GFP-KKT14N/KKT14C from trypanosomes was carried out using anti-GFP antibodies.

### Supplementary Dataset (compressed in one ZIP file)

**Dataset S1**: AlphaFold2-predicted *T. brucei* KKT14^358–685^ structure

**Dataset S2**: AlphaFold2-predicted *T. brucei* KKT14^2–125^ – KKT15 complex

**Dataset S3**: AlphaFold2-predicted *T. brucei* KKT14 – KKT15 complex

**Dataset S4**: AlphaFold2-predicted *T. brucei* KKT14 structure

**Dataset S5**: Files related to the phylogenetic analysis of KKT15/WD40 repeat proteins, including the alignment and corresponding HMM of KKT15 used to collect homologous WD40 families, and folders named ‘Tree1’ and ‘Tree2’, containing the multiple sequence alignments that served as inputs for the phylogenies, and the raw output of the phylogenetic analysis with ‘Tree1’ corresponds to Figure 4, Supplementary Figure S3A, and ‘Tree2’ to Supplementary Figure S3B.

**Dataset S6**: Input alignments and HMMs of KKT14 and KKT15 used for online HHpred.

**Figure S1.**
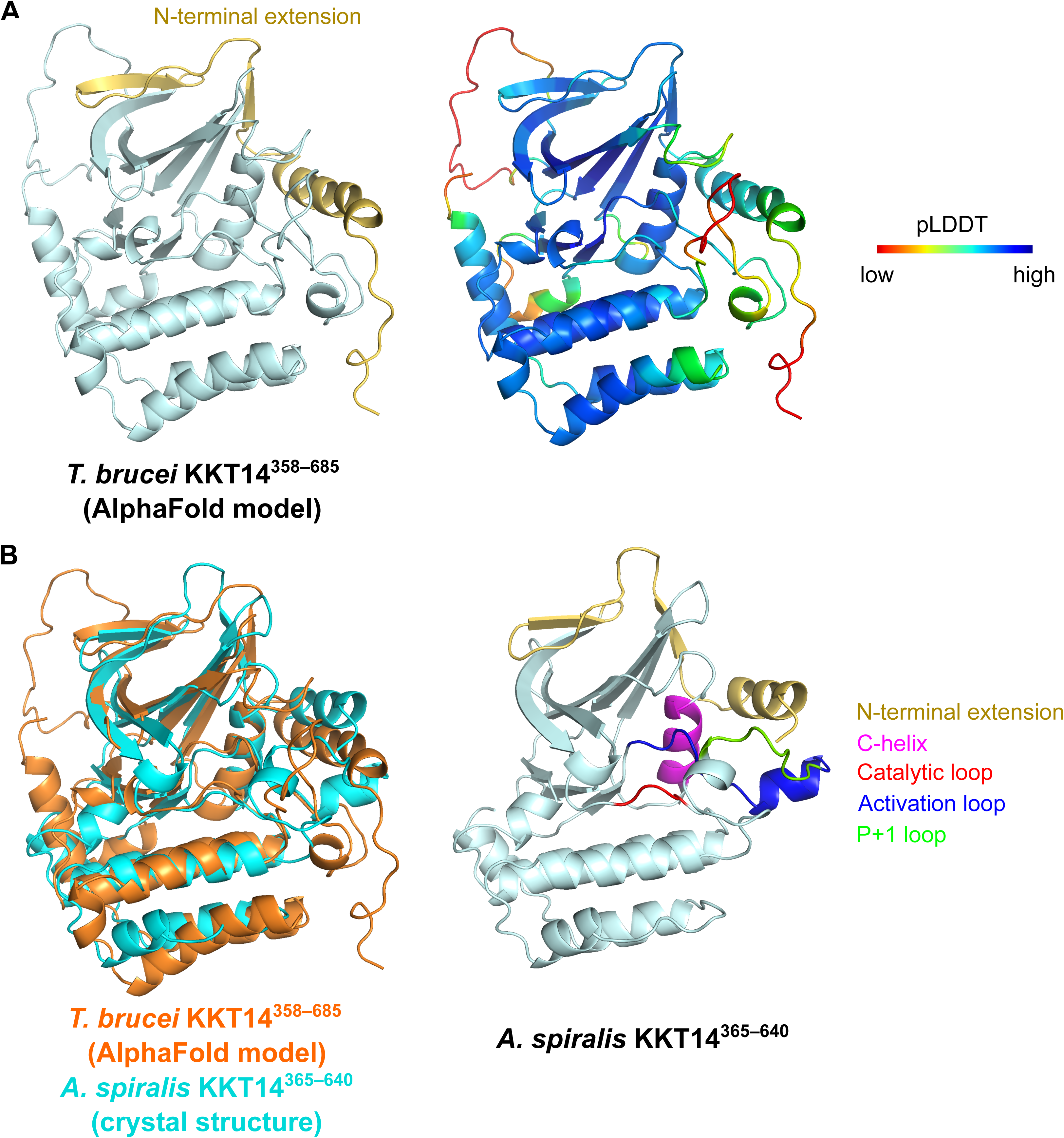
C-terminal pseudokinase domain of KKT14 is conserved in *T. brucei*. (A) Cartoon representation of an AlphaFold2-predicted structure of *T. brucei* KKT14^358–685^. See Supplementary Dataset S1 for the model and pLDDT plots. (B) Overlay of AlphaFold2-predicted structure of *T. brucei* KKT14^358–685^ with a crystal structure of *A. spiralis* KKT14 (duplicated from Figure 1A), showing that positions of N-terminal extension, C-helix, catalytic loop, and part of the activation loop are highly conserved between the two structures.

**Figure S2.**
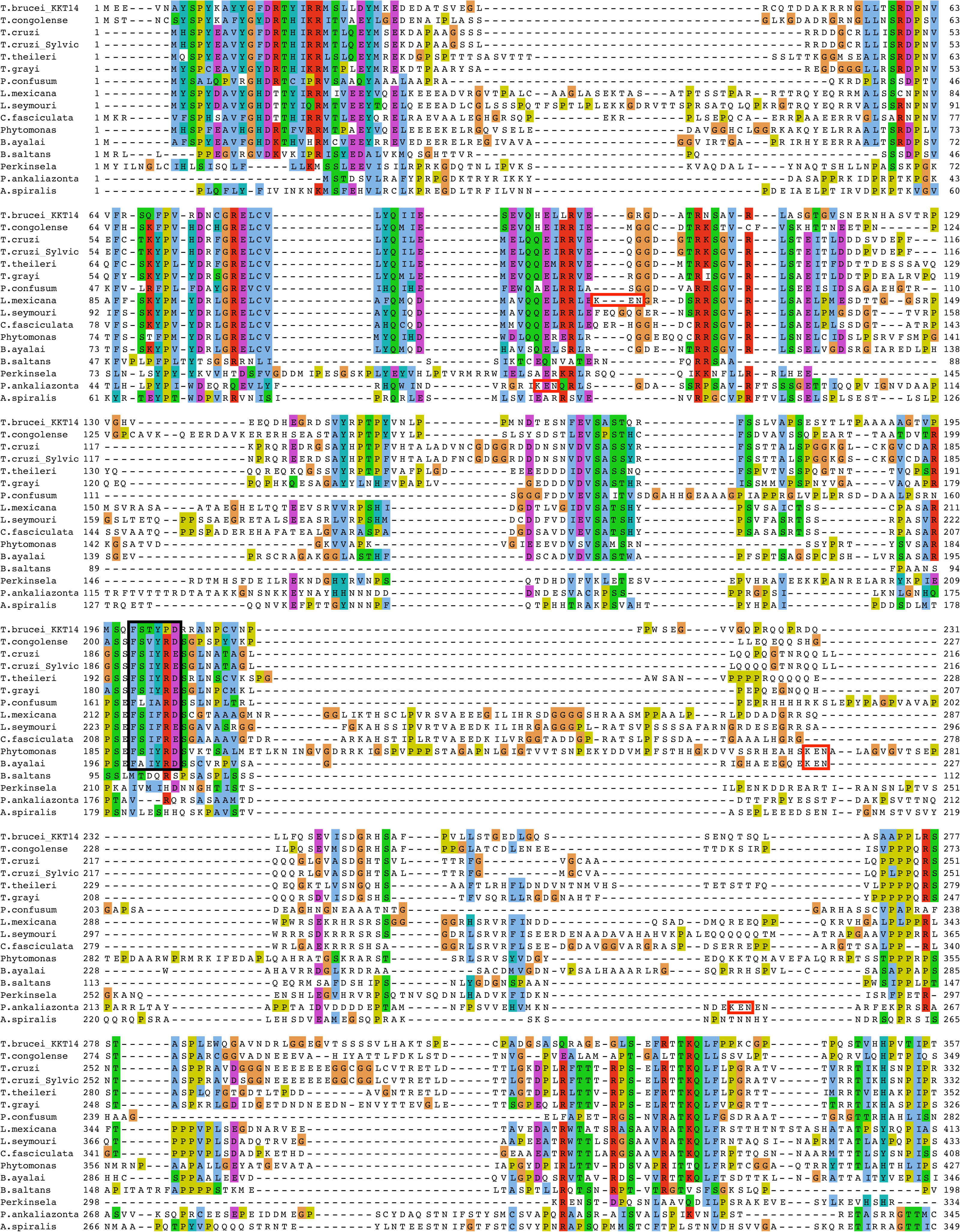
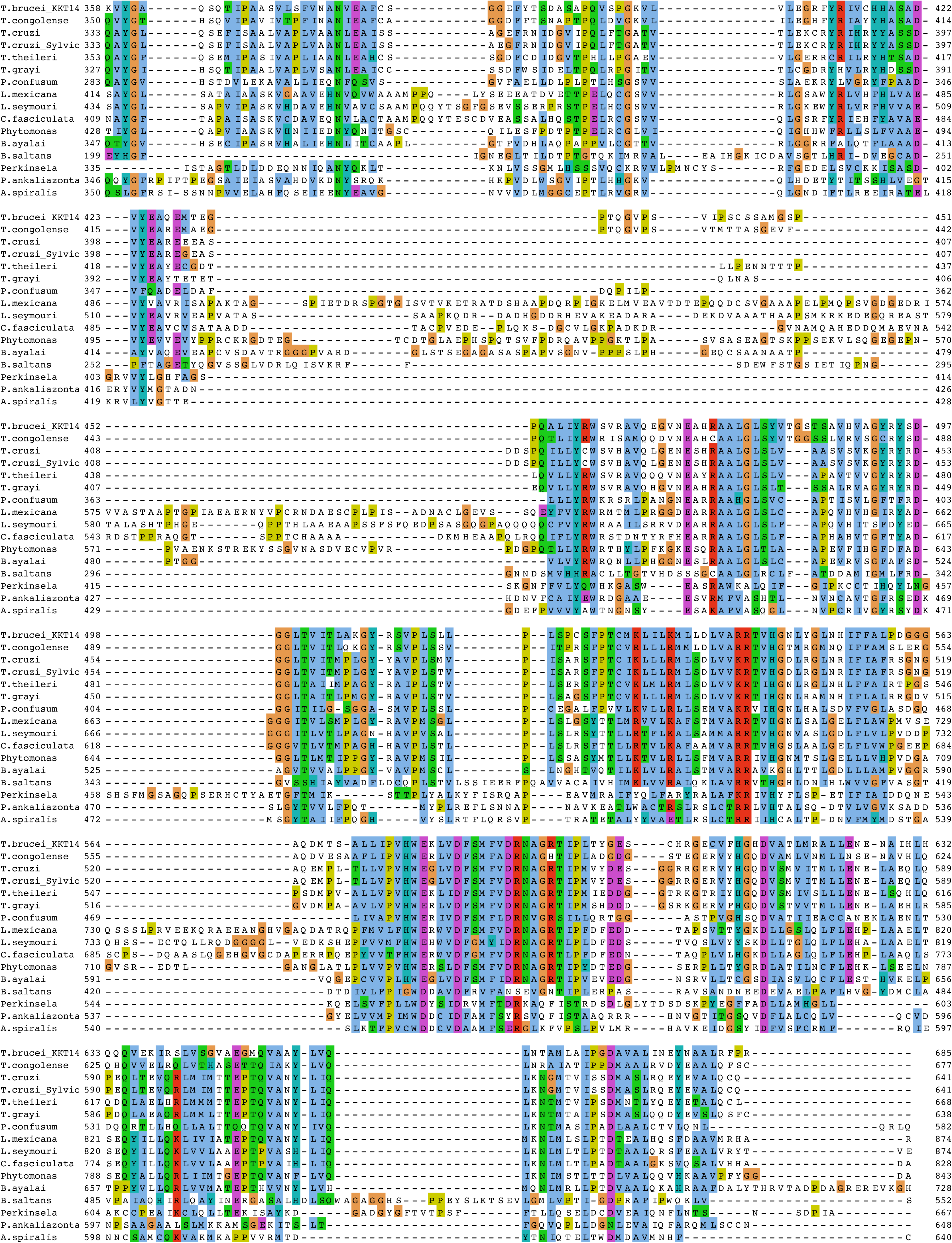
Multiple sequence alignment of KKT14. A putative ABBA motif and KEN boxes are highlighted in black or red boxes, respectively.

**Figure S3.**
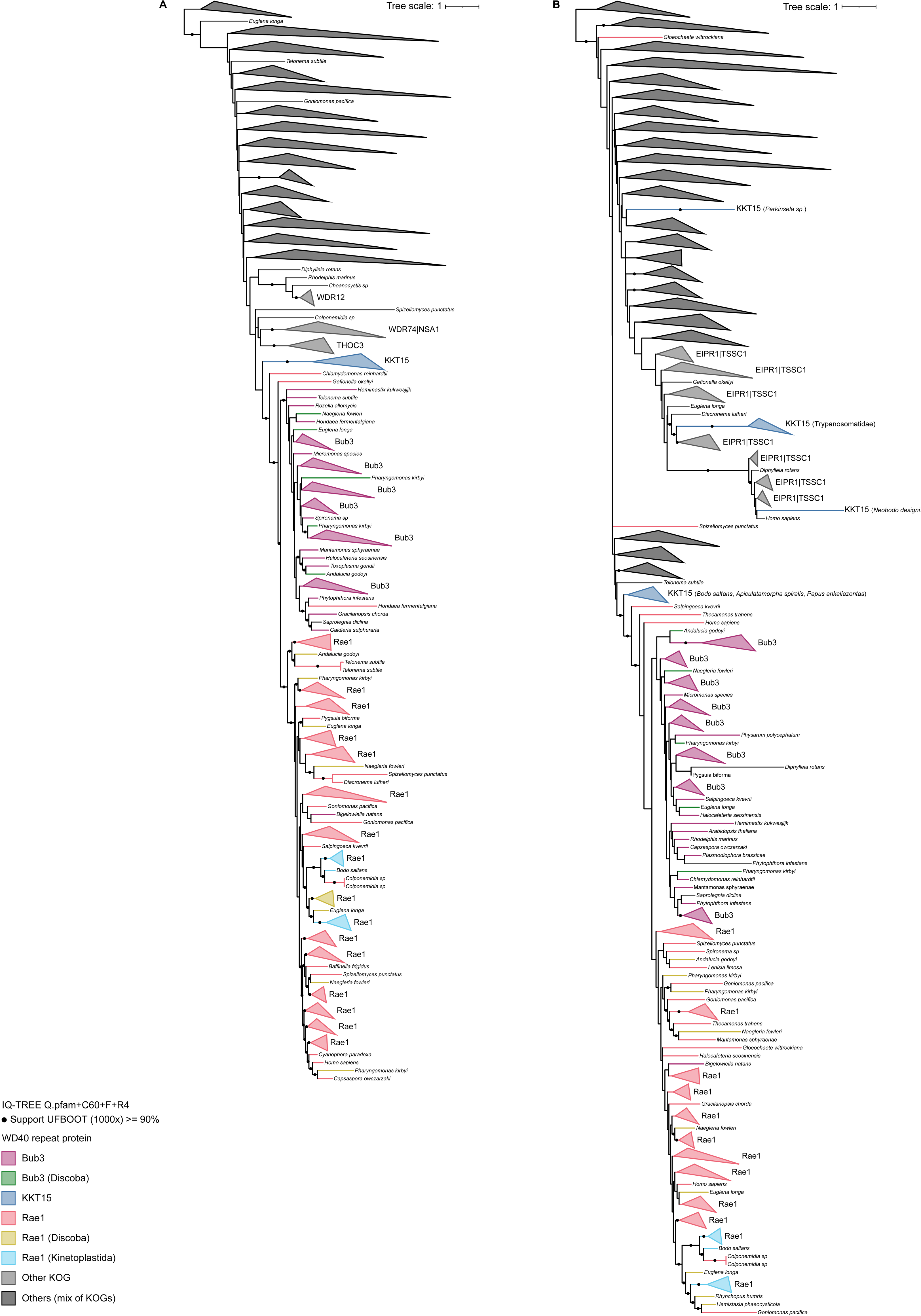
Phylogenies of WD40 repeat proteins, including KKT15. The phylogenies inferred using multiple sequence alignments generated with different approaches. For KKT15, Bub3 and Rae1, and for some potentially related WD40 families (KOG clades), the terminal branches are collapsed into a triangle only if all member leaves correspond to the same group (i.e. monophyletic). Their naming, except KKT15, results from the KOG memberships of the sequences, and was derived from the name of the human member of that KOG as presented in UniProt. If no human member was available for a given KOG, the budding yeast name was used (see Methods). (A) Phylogeny corresponding to Figure 4, of which the alignment approach is described as ‘tree 1’ (Methods). The full, uncollapsed phylogeny can be found here: https://itol.embl.de/tree/62145194227274281702966749 (B) Phylogeny with alignment approach as described under ‘tree 2’ (Methods). The full, uncollapsed phylogeny can be found here: https://itol.embl.de/tree/62145194227421141702968325

**Figure S4.**
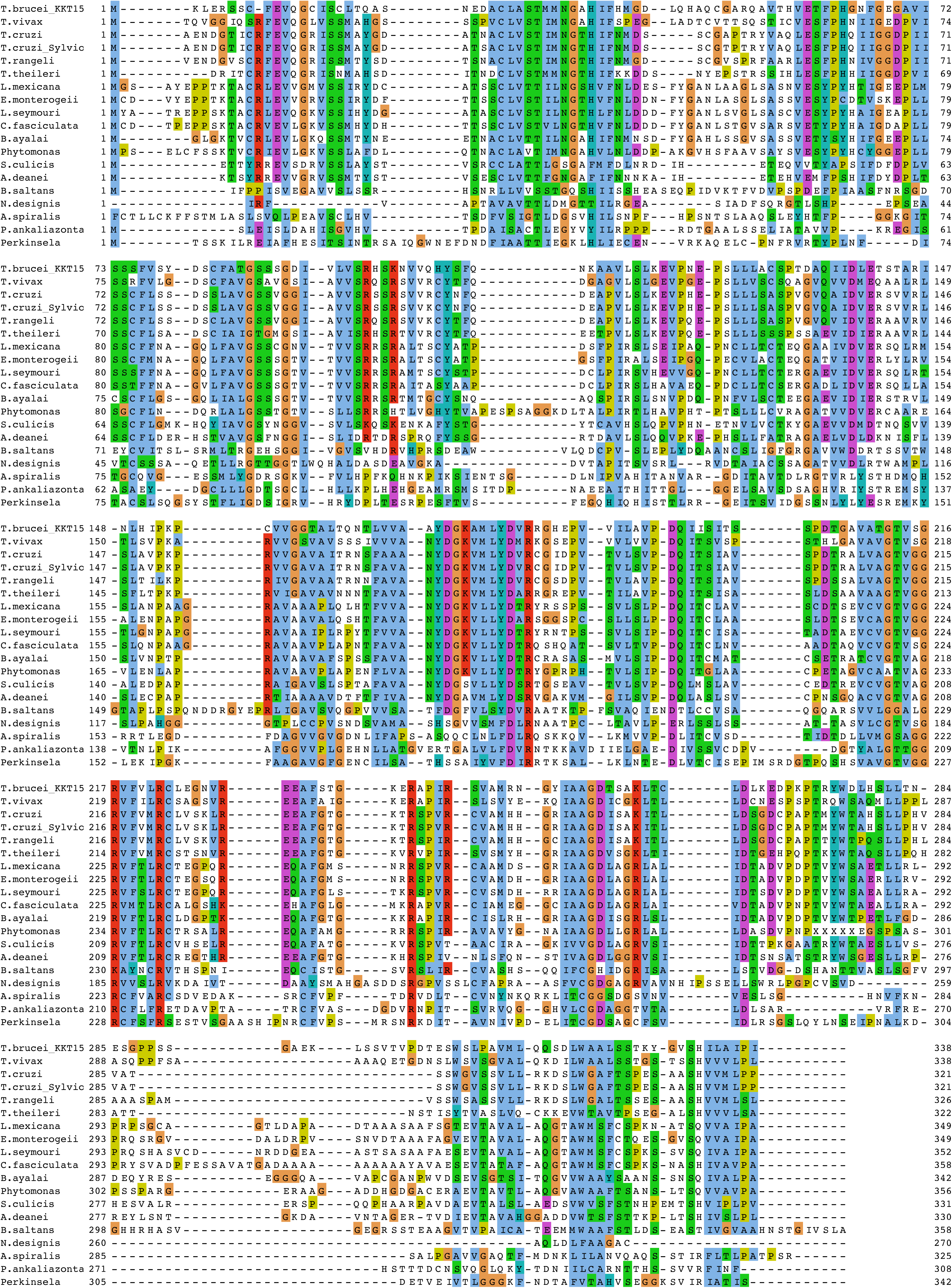
Multiple sequence alignment of KKT15.

**Figure S5.**
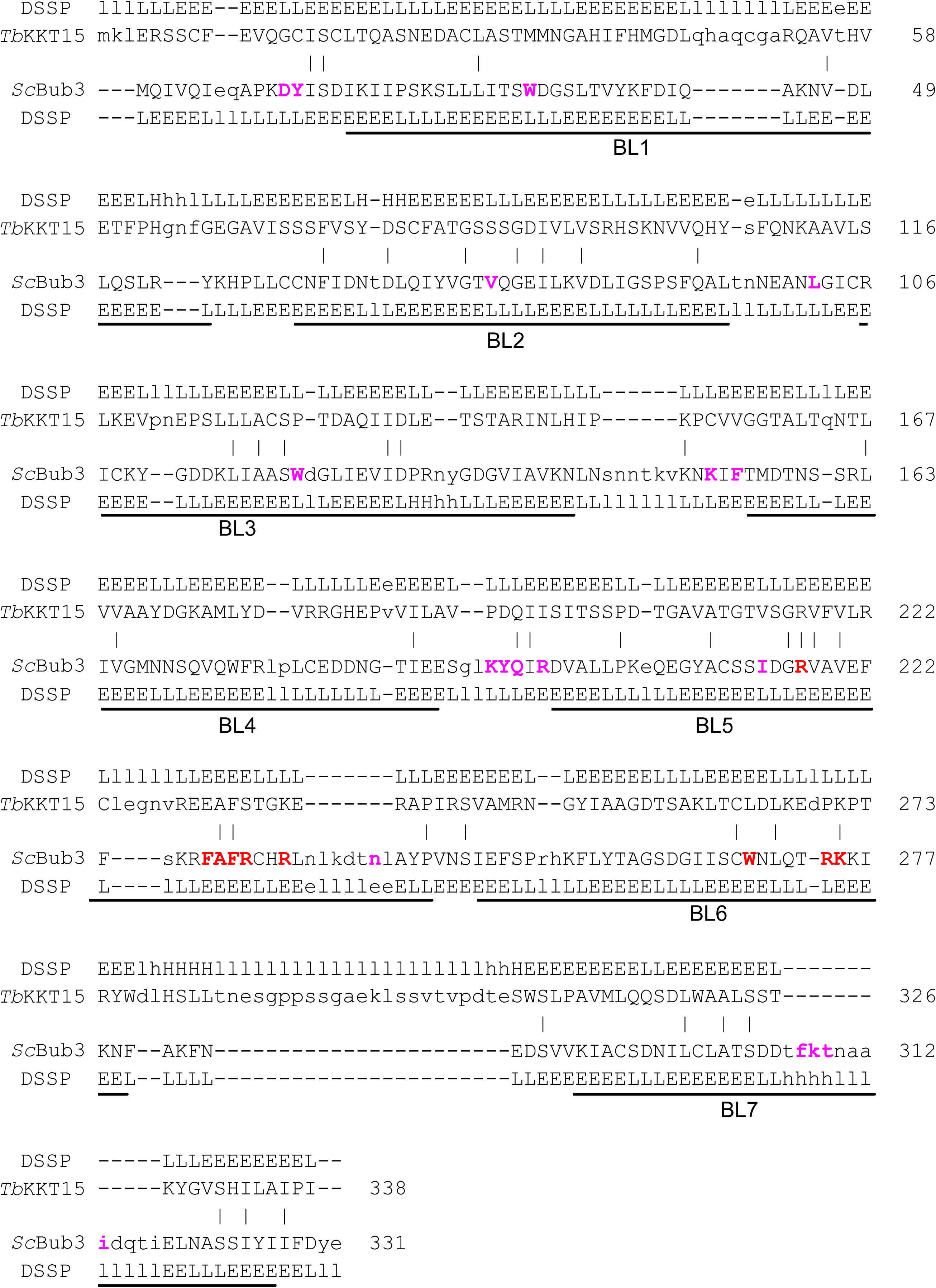
Structure-based pairwise alignment of *T. brucei* KKT15 with *S. cerevisiae* Bub3. DALI search was performed using the KKT15 structure extracted from the AlphaFold2-predicted KKT14^2–125^-KKT15 complex (Supplementary Dataset S2). Structurally equivalent residues are in uppercase, while structurally non-equivalent residues (e.g. in loops) are in lowercase. Amino acid identities are marked by vertical bars. The blade nomenclature of Bub3 (BL1–7) is based on (Primorac et al., 2013). Residues of *S. cerevisiae* Bub3 that interact with Bub1 are highlighted in magenta, while residues that interact with the phosphorylated MELT motif of KNL1 are shown in red as per (Primorac et al., 2013).

