## Supplementary figures and images for "Kinetoplastid kinetochore proteins KKT14-KKT15 are divergent Bub1/BubR1-Bub3 proteins"

### KKT14_24recycles_eeb44_pae.png

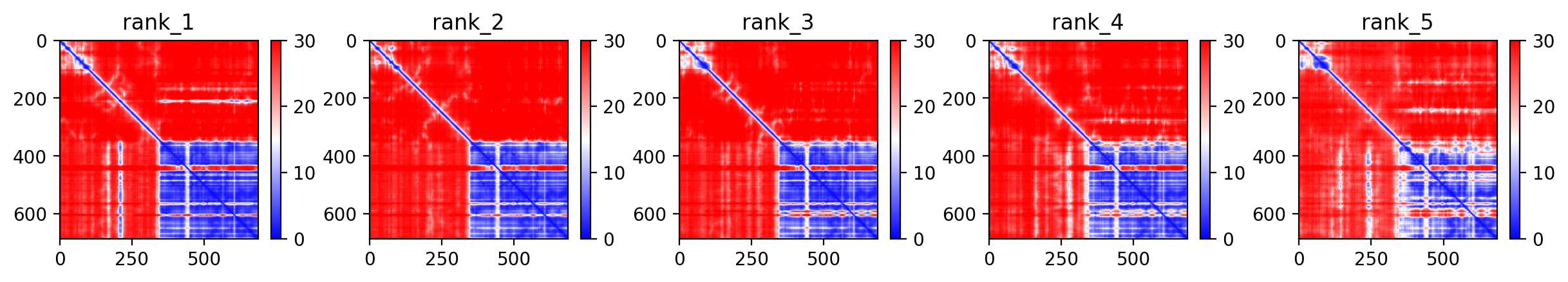

### KKT14_24recycles_eeb44_plddt.png

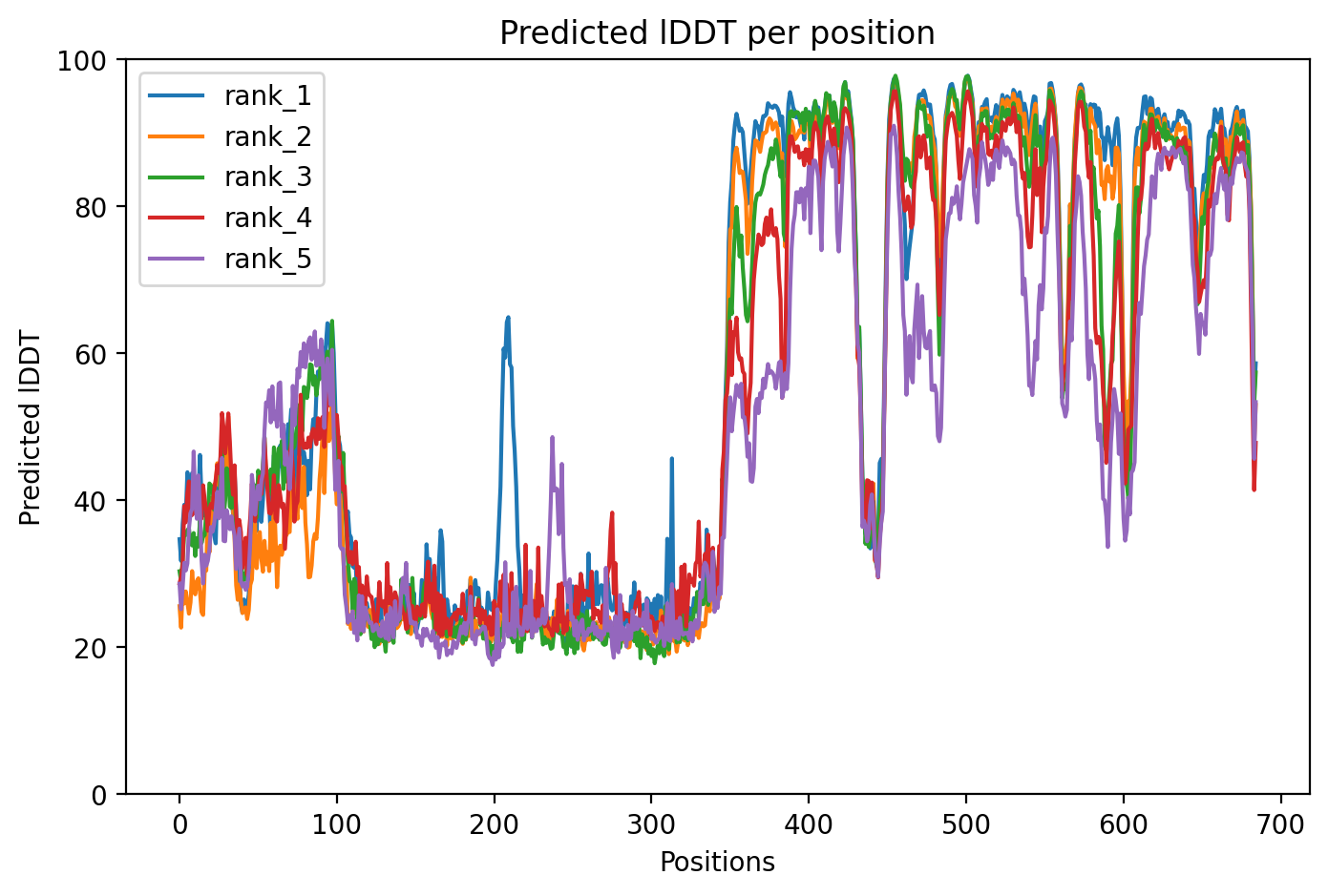

### KKT14_KKT15_24recycles_ad8eb_pae.png

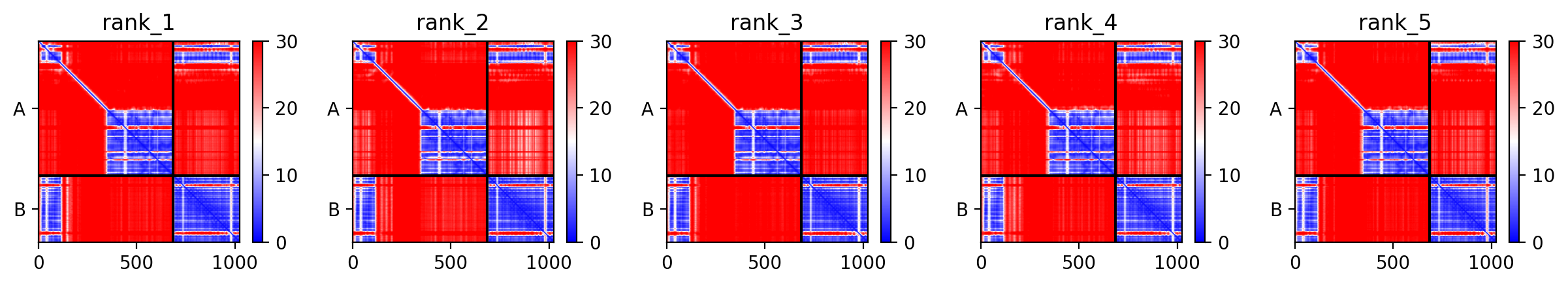

### KKT14_KKT15_24recycles_ad8eb_plddt.png

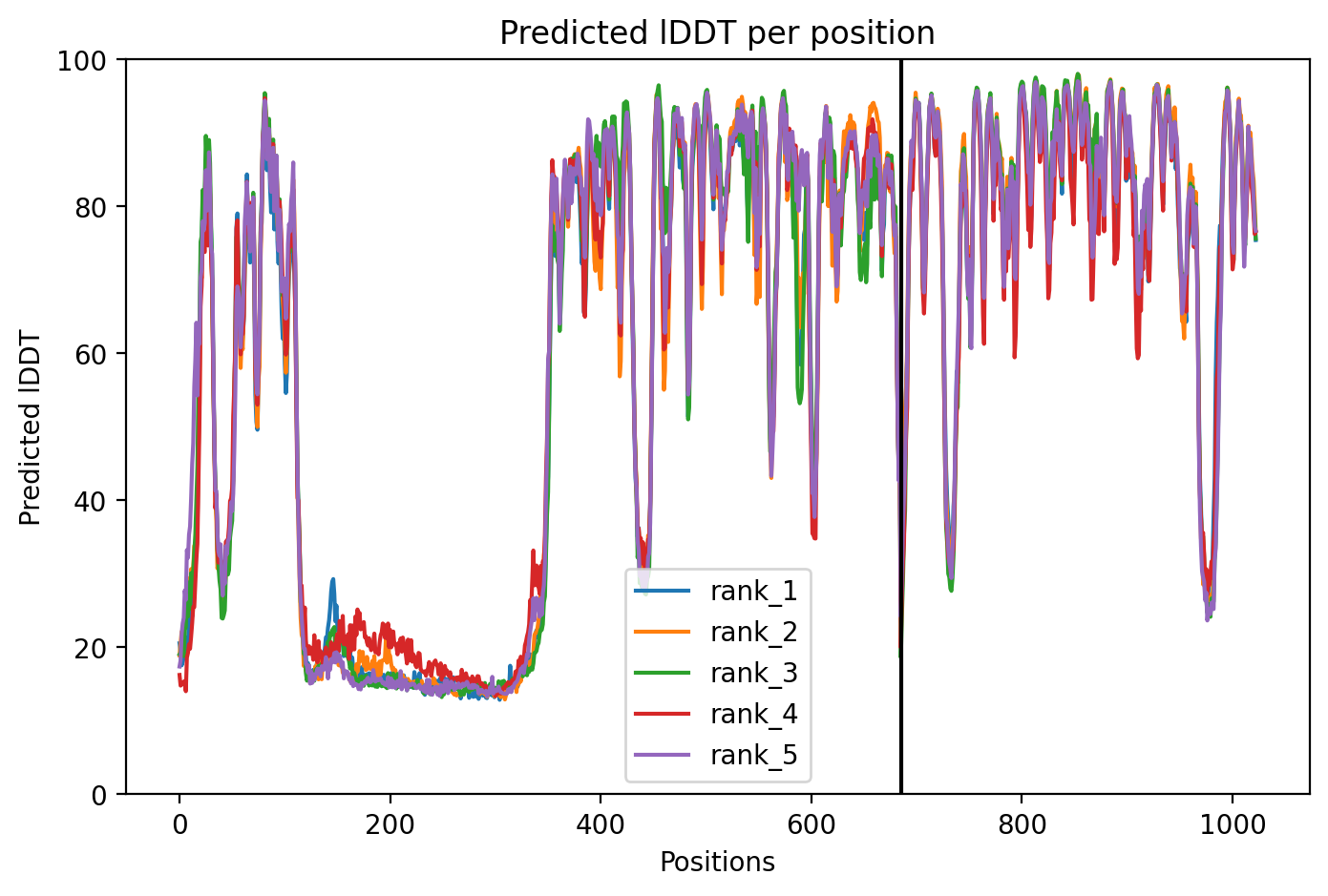

### KKT14C_Tb_24recycles_73e7b_0_pae.png

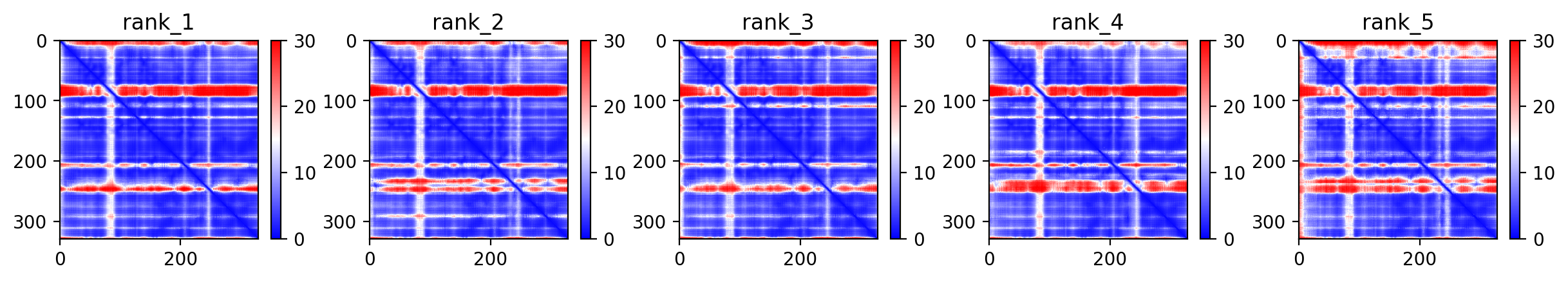

### KKT14C_Tb_24recycles_73e7b_0_plddt.png

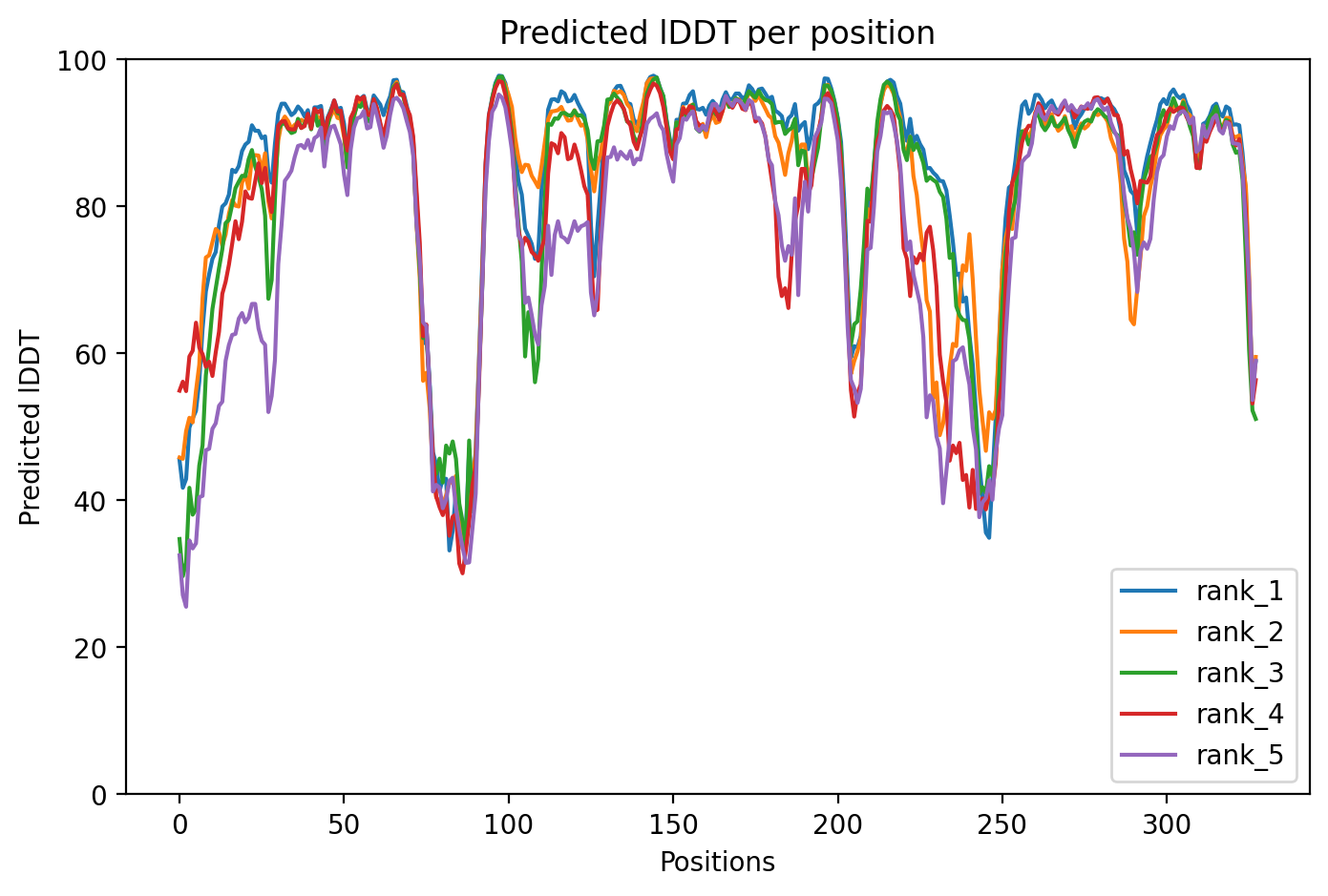

### KKT14N1_KKT15_24recycles_dff6f_pae.png

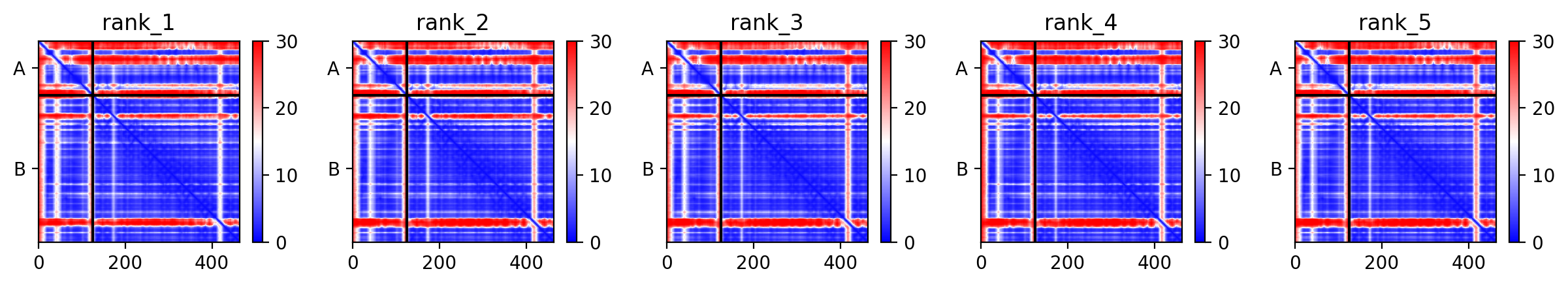

### KKT14N1_KKT15_24recycles_dff6f_plddt.png

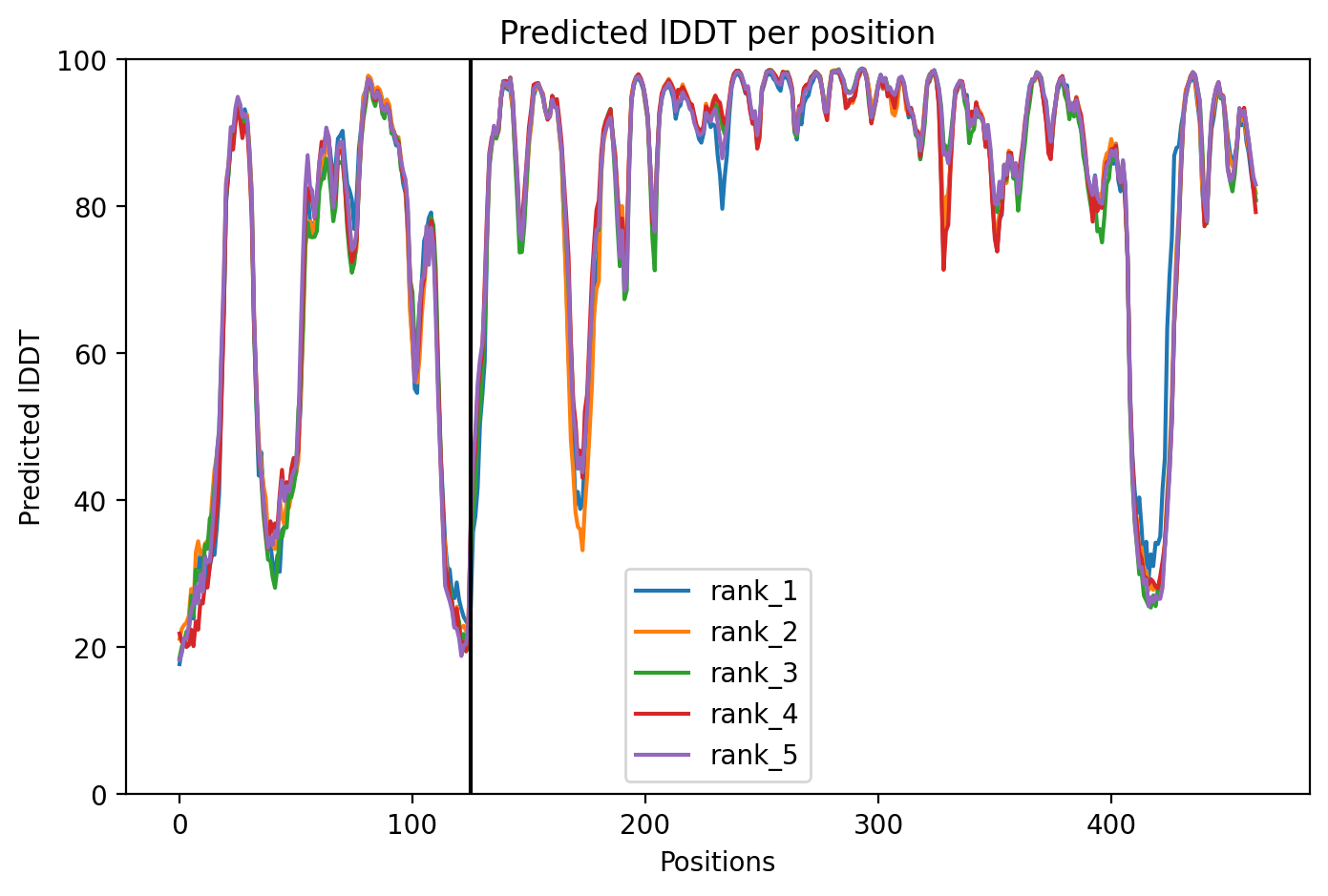
